# Volumetric Compression Shifts Rho GTPase Balance and Induces Mechanobiological Cell State Transition

**DOI:** 10.1101/2023.10.08.561452

**Authors:** Xiangyu Gong, Ryan Nguyen, Zehua Chen, Zhang Wen, Xingjian Zhang, Michael Mak

## Abstract

During development and disease progression, cells are subject to osmotic and mechanical stresses that modulate cell volume, which fundamentally influences cell homeostasis and has been linked to a variety of cellular functions. It is not well understood how the mechanobiological state of cells is programmed by the interplay of intracellular organization and complex extracellular mechanics when stimulated by cell volume modulation. Here, by controlling cell volume via osmotic pressure, we evaluate physical phenotypes (including cell shape, morphodynamics, traction force, and extracellular matrix (ECM) remodeling) and molecular signaling (YAP), and we uncover fundamental transitions in active biophysical states. We demonstrate that volumetric compression shifts the ratiometric balance of Rho GTPase activities, thereby altering mechanosensing and cytoskeletal organization in a reversible manner. Specifically, volumetric compression controls cell spreading, adhesion formation, and YAP nuclear translocation, while maintaining cell contractile activity. Furthermore, we show that on physiologically relevant fibrillar collagen I matrices, which are highly non-elastic, cells exhibit additional modes of cell volume-dependent mechanosensing that are not observable on elastic substrates. Notably, volumetric compression regulates the dynamics of cell-ECM interactions and irreversible ECM remodeling via Rac-directed protrusion dynamics, at both the single-cell level and the multicellular level. Our findings support that cell volume is a master biophysical regulator and reveal its roles in cell mechanical state transition, cell-ECM interactions, and biophysical tissue programming.

## Introduction

Volume regulation is critical in cell behavior, function, differentiation, and survival^1–3^, which are also codependent on the extracellular context^4,5^. The mechanisms underlying the impact of cell volume change on intracellular organization and function and extracellular mechanosensing are not well understood. Moreover, native tissue microenvironments that are composed of a scaffolding extracellular matrix are complex materials with non-elastic properties (e.g. stress relaxation, strain-induced stiffening, alignment, densification, and plasticity)^5,6^. Together, these render a challenging, dynamic picture of cell mechanosensing and biophysical phenotype regulation in the body.

The cytoskeleton provides mechanical support and enables physical functions, such as migration and force generation. It is a highly dynamic network of filaments and accessory protein complexes that are constantly undergoing turnover and interactions. Its organization and spatiotemporal profile dictate mechanical outputs and physical interactions with the cell’s surroundings. Actin filaments are the key base components, involved in cell protrusions (lamellipodia, filopodia), force generation (stress fibers, actin cortex), and structure^7,8^. Rho GTPases, notably Rac1, RhoA, and CDC42 are upstream signals that control actin dynamics and cytoskeletal tension through modulating actin nucleation factors and myosin II^9,10^. These ultimately determine how the actin cytoskeleton is organized and programmed.

The extracellular matrix that scaffolds cells in most tissue microenvironments is a complex network of proteins. Collagen I is one of the most prominent components and provides the fibrillar architecture found in many tissues. Unlike standard elastic or purely rigid substrates used in many mechanosensing studies, the ECM is a complex material with non-elastic properties. ECM fibers are interconnected through various bonds that can unbind or become rearranged under applied forces, which can lead to stress relaxation and permanent restructuring of the local architecture and ligand distribution^6,11,12^. Non-elasticity confers mechanics-driven time-dependence in ECM properties. The interfacing of the dynamic cytoskeleton with the non-elastic ECM through physical contact and adhesions results in spontaneous physical interactions together with biophysical signaling and feedback that underlie tissue development, homeostasis, and pathology^5,13^. While the functions of various cytoskeletal features are well known on stiff or elastic substrates, their functions in non-elastic microenvironments are much less established.

Cell volume changes can be induced by various biophysical stimuli, including osmotic pressure, mechanical compression, and alterations in cell spreading (e.g. via changes in substrate stiffness)^1,3,14^. Hyper-osmotic pressure and mechanical compression can both reduce cell volume through water efflux, resulting in increased intracellular crowding, cytoskeletal concentration, and substantial changes to the physical profile of the cell^3,14^. Volume modulation can occur both physiologically and pathologically^1,2^. However, it is not well understood how it transforms the cytoskeletal and mechanosensing machinery and biophysical cell and tissue phenotypes, especially in physiologically relevant non-elastic microenvironments.

Here, we investigate the role of volumetric compression in cytoskeletal reprogramming and its impact on mechanosensing and cell-matrix interactions. We compress cells via osmotic pressure from water soluble, low molecular weight polyethylene glycol (PEG300)^3,14,15^. We reveal that Rho GTPase signaling is altered by volumetric compression. A ratiometric shift between active Rho and Rac alters cytoskeletal organization and cell shape dynamics; stress fibers across the cell interior and fluctuating lamellipodia are reduced while peripheral actin is increased along with the presence of altered cell adhesions. This modulates mechanosensing, particularly on non-elastic collagen I substrates, which is dependent on dynamic cell boundary fluctuations. Furthermore, we demonstrate that a rebalancing of Rho GTPase activities can override the cytoskeleton state under compression and partially rescue the non-compressed cell phenotypes.

## Results

### Sustained restriction in cell volume alters cell spreading behavior and morphodynamics

In biological processes, such as cancer and development, or after surgical procedures, such as implantation or injection, cells are subjected to mechanical compression in a complex microenvironment with cell-cell and cell-matrix interactions (Fig. 1A). For instance, in hepatocellular carcinoma, we found some regions of the tumors exhibit a gradient of nuclei size and density from the stroma toward the inner region of the tumor (Fig. 1B, Patient 1). Furthermore, in regions where a tumor grew inside vasculature (microvascular invasion), we observed both compressed endothelial cells and more packed, smaller tumor cells between the solid tumor and the stroma (Fig. 1B, Patient 2). Abundant studies have supported that dysregulated growth of tumors induces compression on both constitutive cancer cells as well as the normal adjacent cells^16–18^, which possibly leads to volumetric compression on cells^19^ and changes in cell shape and behaviour^20^. Here, we introduce volumetric compression on mesenchymal-like liver cancer cells (SNU-475) via polyethylene glycol (PEG)-induced colloidal osmotic pressure and allow for cell spreading on a collagen-coated tissue culture plastic surface. After 24 hours of cell spreading, we found that the PEG-induced compression leads to a reduction in cell spreading area and controls cell morphology (Fig. 1C). A higher PEG concentration results in a smaller cell spreading area. These compressed cells also adopted a more “solid-like”, convex shape with less irregularity, characterized by the metric “solidity” (Fig. 1D). To visualize cell shape and volume in 3D, we loaded the cells with CellTracker, a live cytosolic dye, before plating the cells to attach under compression for 24 hours (Fig. 1E). Consistent with the previous studies^3,14,21^, the inert soluble polymer PEG-300 in culture media persistently induces compression on cells to reduce cell volume (Fig. 1F,G) and subsequently condenses and increases concentrations in the cytosol indicated by the fluorescent intensity of the CellTracker signal (Fig. 1H).

**Figure 1.**
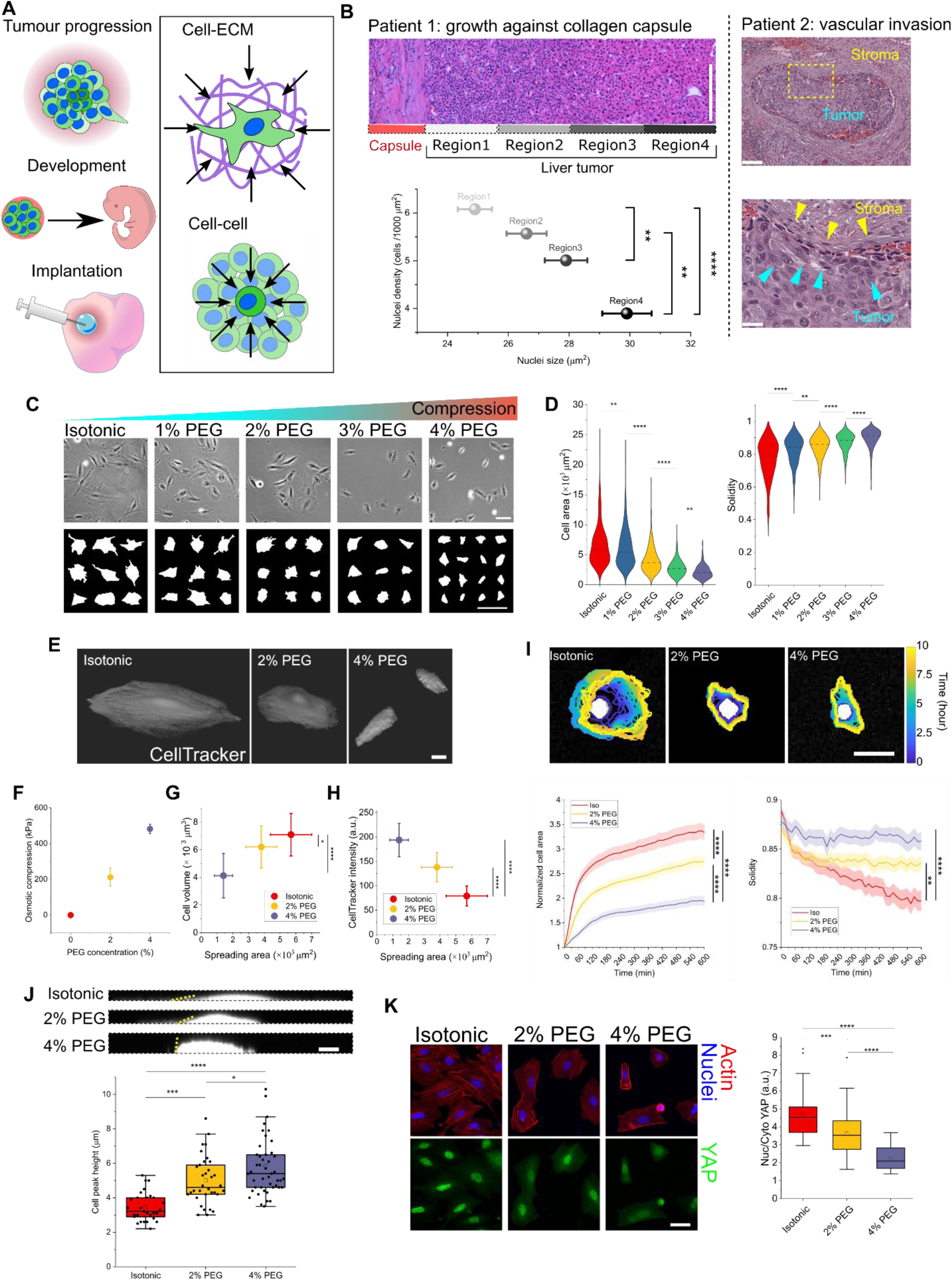
Osmotic compression induces a persistent reduction in cell volume and affects cell spreading dynamics. (A) Schematic illustrating examples of compression-involved biological processes where cells interact with each other or with the surrounding ECM. (B) HCC histological samples exhibit gradients of cell size and density near the tumor-fibrotic capsule interface (Patient 1) and highly compressed regions of cells due to tumor expansion during microvascular invasion (Patient 2). Yellow arrows: stromal cells; blue arrows: tumor cells. Scale bars, 200 μm. (C) Brightfield images and cell shape segmentation showing 2D cell morphology is a function of intracellular osmotic pressure, corresponding to the measurements of cell area and solidity in (D). Every condition is significantly different from the others. N=339-371 cells per condition. Scale bar, 200 μm. (E) 3D rendering of CellTracker signal visualizing cell morphology under different osmotic conditions. Scale bar, 15 μm. (F) Measurements of osmotic pressures generated by isotonic (Iso) culture medium or varying concentrations (compression media: 2% and 4%) of PEG-300. n: 4-5 measurements per condition. (G) Measurements of cell area and cell volume under different levels of compression after 24-hour cell attachment. (H) Measurements of cell area and intracellular mean intensity of CellTracker under different levels of compression after 24-hour cell attachment. For (G) and (H): N=29 cells for isotonic, 34 cells for 2% PEG, and 46 cells for 4% PEG. (I) Temporal tracking (10 hours, time interval: 15 min, Supplementary Video 1) of cell area upon seeding of the CellTracker-labled cells in compression media (Iso, 2% PEG, 4% PEG). Respectively, N=89, 82, and 84 cells. Scale bar, 80 μm. (J) Profiling cell heights under the conditions of Iso, 2% PEG, and 4% PEG after 24-hour cell attachment. Respectively, N=29, 34, and 46 cells. Scale bar, 10 μm. Yellow dotted lines indicate the contact angle of the cells with respect to the substrate. (K) Representative fluorescent images of YAP and F-actin, and YAP nuclear translocation under varied compressions. N=31, 32, and 31 cells respectively. Scale bar, 50 μm. Significant comparisons were determined with a one-way ANOVA with a Tukey post hoc test. (*p<0.05; **p<0.01; ***p<0.001; ****p<0.0001). All experiments were conducted three times independently. The lines in the violin plots represent median values. The boxes in the box blots represent the 25%-75% interquartile range (IQR) and whiskers represent 1.5IQR. In (B), (F), (G), (H), data represent mean±s.d. In (I), data represent mean±s.e.m.

Cells in the mesenchymal state are highly dynamic and fluid-like to enable morphogenesis^22^ and cancer metastasis^23^. Studies have shown that under mechanical stress, mesenchymal cells undergo a dramatic transition in spreading state and migration mode^24^. We investigate how volumetrically compressed mesenchymal cancer cells adapt their spreading state on the rigid 2D surface dynamically (Supplementary Video 1). Ten-hour time-lapse imaging reveals that under compression the cells spread into smaller areas at a lower rate (Supplementary Fig. 1A) with much less volatile edges (Fig. 1I). This is reflected by the soon plateaued cell solidity under compression, while cells in the isotonic condition have rapidly changing cell shape with an increasing irregularity, indicated by a continuously decreasing solidity. When fitting the cell centroids’ movement during cell spreading into the anomalous diffusion model^25,26^, we observed less persistence, more confined, and subdiffusive motion under compression (Supplementary Fig. 1B-F). Furthermore, after 24 hours of cell spreading, the compressed cells form “bulge-like” shapes, whose heights and contact angles are significantly higher than the “sheet-like” non-compressed cells (Fig. 1J). This indicates that volumetric compression limits cell spreading capability. Literature has shown that cell spreading, cytoskeleton, tension, and nuclear deformation dictate subcellular translocation of YAP/TAZ, a master regulator of mechanostransduction^27–29^. We find the compressed cells with restricted cell spreading indeed have reduced YAP nuclear localization and lack perinuclear stress fibers (Fig. 1K). After fully spreading (~ 24 hours), the cells in the isotonic condition exhibit dramatic morphodynamics with ruffling and cell area fluctuation over two hours, while compressed cells show significantly reduced shape fluctuation and a rather stable cell morphology and area (Fig. 2A,B and Supplementary Fig. 2A, Supplementary Video 2).

**Figure 2.**
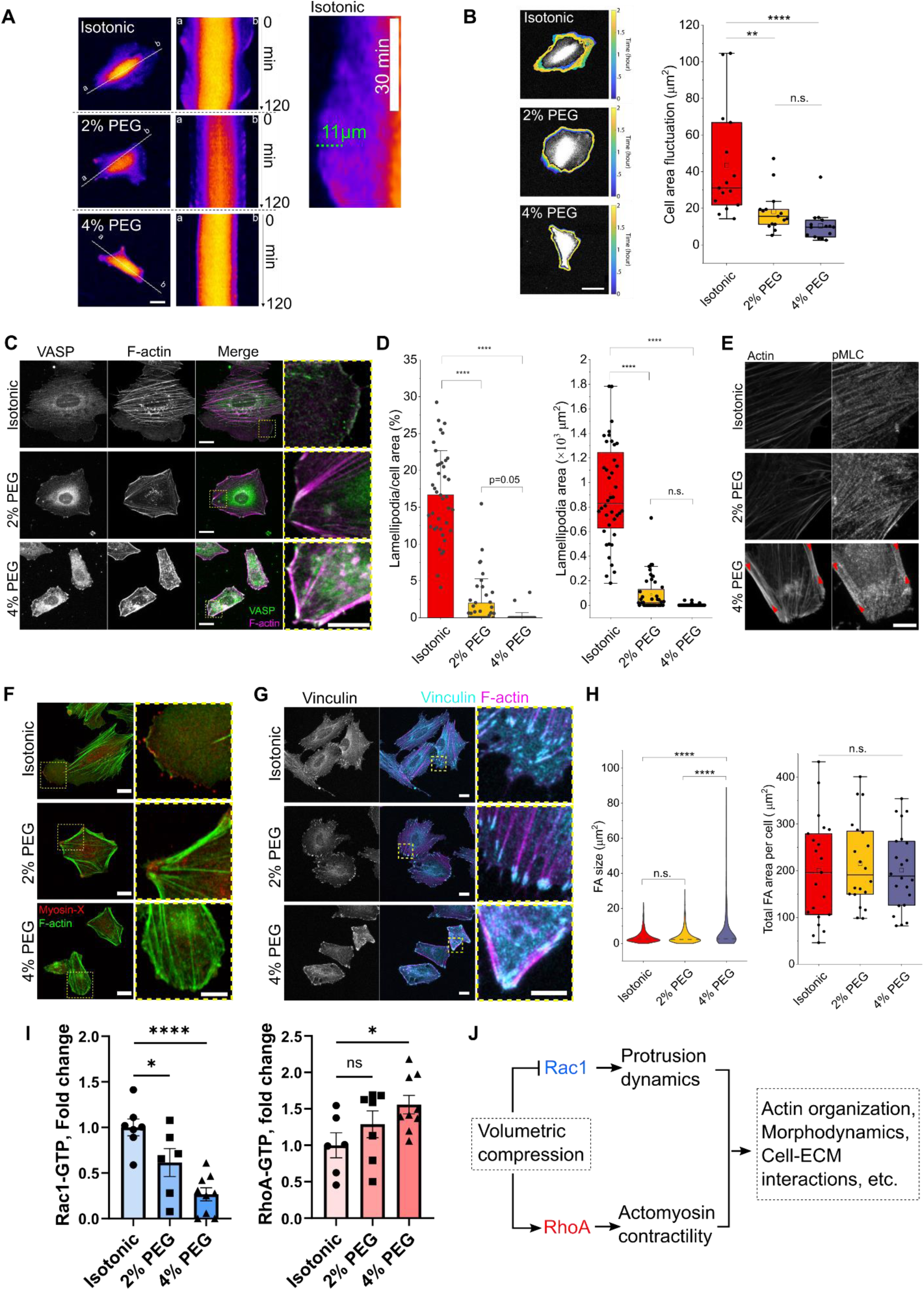
Cell morphodynamics and adhesion-actin machinery respond to volume-regulated Rho/Rac balancing. (A) Kymographs showing 120-min cell edge ruffling after 24-hour cell attachment in compression media. Scale bar, 25 μm. (B) Profiling cell boundaries for 120 min (2 min interval) and measuring area fluctuation of individual cells, defined by the standard deviation of the mean area of the cell over time (also see Extended Data Fig.1A, Supplementary Video 2). N=15 cells per condition from three independent experiments. Scale bar, 50 μm. (C) Fluorescent images of VASP and F-actin of the cells under varied compression. The close-up images show the distribution of VASP with respect to the actin. The fluorescent images were used to characterize: (D) the ratio of lamellopodia in the whole cell area and the absolute area of the lamellopodia. N=40, 40, and 52 cells respectively from three independent experiments. (E) High-resolution confocal images show the relative distribution of p-MLC on the actin cytoskeleton. (F) Fluorescent images of Myosin-X and F-actin of the cells under varied compression. Scale bar, 10 μm. (G) Fluorescent images of Vinculin and F-actin of the cells under varied compression. The close-up images show the distribution of vinculin with respect to the actin. (H) Vinculin puncta were used to determine the size distribution of focal adhesions and the total adhesion area of individual cells for varied conditions. N=21, 20, and 22 cells respectively from two independent experiments. FA measurements from all cells were pooled to plot the size distribution. (I) Quantifications (GLISA) of active (GTP-bound) RhoA and Rac1 of the cells cultured under varied compression for 24 hours compared to the cells culture in the isotonic condition. Mean±s.e.m. Statistic differences were calculated by one-way ANOVA with a Dunnett’s multiple comparisons test. (J) Proposed model of volumetric compression regulating Rho/Rac balance. For (C,E,F), scale bars in the close-up images, 20 μm; scale bars in the close-up images, 10 μm. Statistic analyses in (B,D,G) were calculated by one-way ANOVA with a Tukey multiple comparisons test (*p<0.05; **p<0.01; ***p<0.001; ****p<0.0001; ns: not significant).

### Volumetric compression regulates the actomyosin-adhesion machinery via Rho and Rac re-balancing

Actin cytoskeletal arrangement is a direct determinant of cell morphology, force generation states, migration, cell-cell communication, and cell interactions with the microenvironment. To understand the impact of volumetric compression on cell spreading dynamics, we investigate actin cytoskeleton and focal adhesion using immunofluorescence staining after 24 hours of cell spreading. We observe distinct actin arrangements under varied volumetric compression (Fig. 2C-E). The “solid-like” morphology and the limited dynamics of compressed cells are associated with (1) the formation of thick actin bundles on the cell periphery that are rich in phosphorylated myosin light chain (pMLC) (Fig. 2E, Supplementary Fig. 2D), and (2) the loss of VASP-rich lamellipodia. Further, abundant VASP is accumulated at the adhesion sites of the peripheral actomyosin bundles (Fig. 2C, Supplementary Fig. 2B), which has been associated with tension maintenance in confluent epithelium^30^. In the isotonic condition, the lamellipodia of cells made up approximately 15% of the cell area on average, whereas cells have significant area reduction in the lamellipodia portion under 2% PEG compression and almost no lamellipodia under 4% PEG compression (Fig. 2D). The impeded growth of lamellipodia subsequently results in the loss of Myosin-X directed filopodia (Fig. 2F). Vinculin, a component of the focal adhesion (FA) complex, also exhibits altered morphology and distribution (Fig. 2G, Supplementary Fig. 2C). In the isotonic condition, the cells can form nascent adhesions to guide lamellipodium growth, which are absent in the compressed cells. However, in all conditions, cells formed mature FA plaques to anchor and establish actin stress fibers or peripheral actomyosin bundles. Interestingly, the total FA area per cell is conserved despite the reduction in cell area under the compressed conditions (Fig. 2H, Supplementary Fig. 2C), which give rise to fewer but larger vinculin plaques in highly compressed (4% PEG) cells. The conserved total FA area suggests a conserved anchorage force for the adhering cells regardless of the volumetric compression^31,32^.

Small GTPases Rho and Rac are master regulators of the actin cytoskeleton. Rho-associated kinase (ROCK), a key downstream effector of RhoA, phosphorylates myosin to generate contractile force, whereas Rac1 facilitates actin elongation, protrusion, and lamellipodia formation^33^. The intricate balance between Rho and Rac determines cell morphology, migratory phenotypes, and cell functions^34–37^. Previous studies have shown physical confinements or hyperosmolarity triggers actin cytoskeletal response mediated by Rho/ROCK ^24,38–40^. Here, without compression, Rho and Rac in the mesenchymal-like SNU475 cells coordinate in such a way that the cells spread into thin sheets with pronounced internal actomyosin stress fibers and dynamic lamellipodia. We hypothesize that under volumetric compression, the shifted cell morphology with the assembly of actomyosin bundles on the cell periphery and the loss of lamellipodium is linked to a volume-regulated Rho/Rac rebalancing (Fig. 2J). This is confirmed by measuring active (GTP-bound) RhoA and active Rac1 from the same batches of cells in the compression conditions after 24 hours of spreading (Fig. 2I). We find that under high volume compression (4% PEG), cells have elevated Rho activation and significantly reduced active Rac1. Interestingly, 2% PEG compression did not render a detectable enhancement in RhoA activity but was sufficient to suppress Rac1 activity. This compression-suppressed Rac1 activity explains the loss of lamellipodia and reduced morphodynamics. To evaluate the Rho-related actomyosin machinery in the compressed cells, we measure the densities of the actin cytoskeleton and pMLC from fluorescent images (Supplementary Fig. 2D-G). Notably, compared to the isotonic case, the mid-level compressed (2% PEG) cells have a higher density of pMLC but a similar pMLC density colocalized with F-actin (pMLC intensity/F-actin intensity). In the highly compressed (4% PEG) cells, although F-actin and pMLC are both highly condensed, the normalized pMLC by the total actin is decreased (Supplementary Fig. 2G), suggesting a lower level of contractile stress despite a high Rho activation. This supports the prior finding that both Rho activity and the extent of cell spreading impact myosin phosphorylation^35^.

### Volumetric compression dictates elastic and non-elastic cell-substrate interactions

To further investigate cellular force and how cells physically interact with their surroundings under volumetric compression, we evaluate cell contractility using both traction force microscopy (TFM) and the collagen compaction assay (Fig. 3). Based on methods developed previously^41–44^, we characterized cell traction force by computing the strain energy generated on elastic polyacrylamide (PA) substrates with a ~3 kPa shear storage modulus (Supplementary Fig. 3A). Consistent with the culture on a rigid substrate, under compression attaching cells adopted a “solid-like” morphology (Fig. 3A) and a reduced spreading area (Fig. 3B). To our surprise, strain energy generated by the cells is conserved under 2% PEG compression compared to isotonic (Fig. 3C) despite a smaller cell area, consistent with a higher strain energy density of these compressed cells (Fig. 3D). Under the higher compression (4% PEG), the cells no longer generate the same level of strain energy, yet maintain a comparable level of strain energy density with a much smaller cell spreading area. The results of TFM imply that volumetric compression does not suppress or inhibit force generation machinery (energy density) but can influence the total mechanical output (strain energy) based on the cell spreading area.

**Figure 3.**
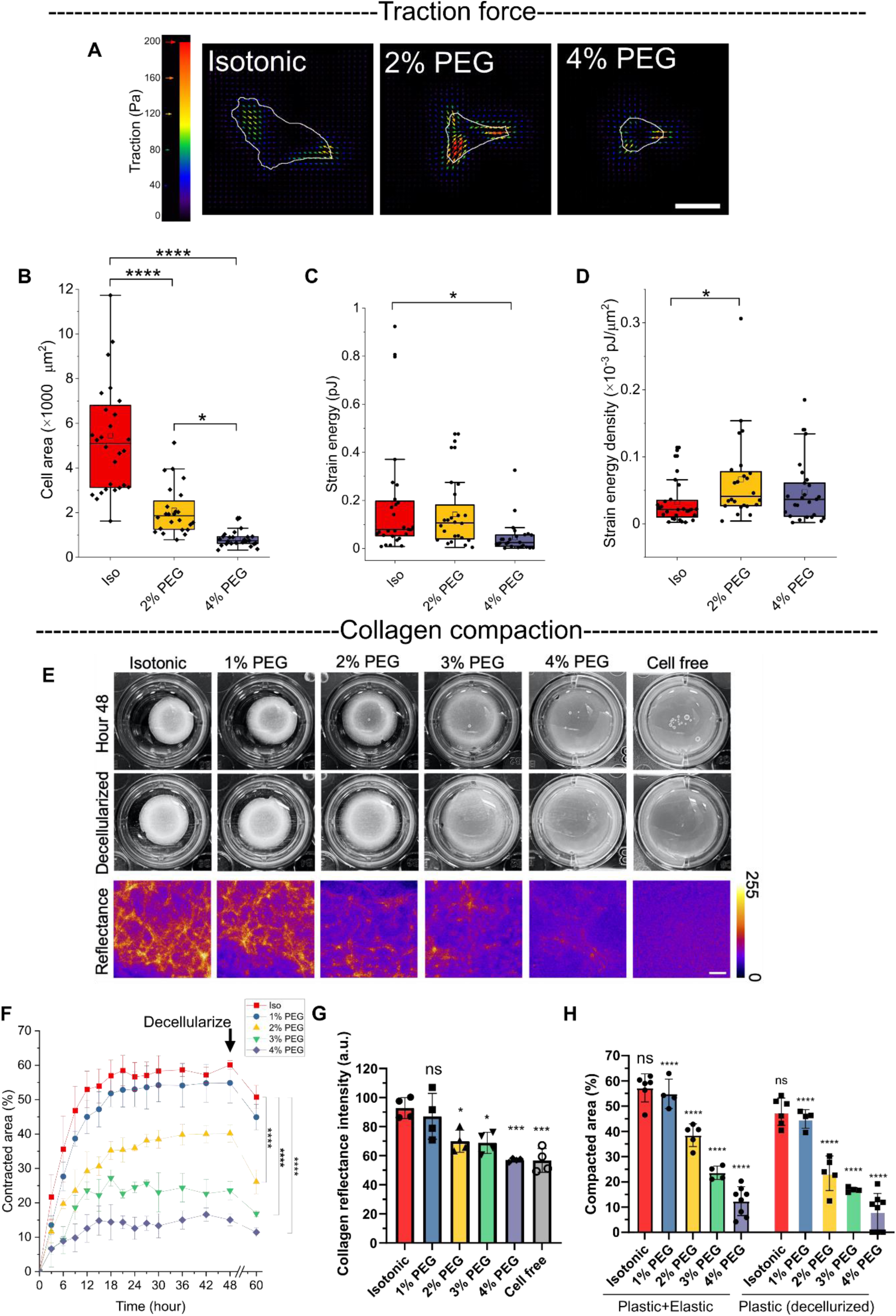
Actomyosin contractility remains intact, but cell-ECM interactions are altered under compression. (A) Representative stress-field maps of TFM of cells under isotonic and compression conditions. Scale bar, 50 μm. Cell area (B), calculated cell strain energy (C), and strain energy density (D), were measured from TFM. N=28, 24, and 25 cells per condition from two independent experiments. The boxes in the box blots represent the 25%-75% interquartile range (IQR) and whiskers represent 1.5IQR. (E) Collagen contraction assay showing contracted collagen area before and after decellularization under different levels of compression, and the corresponding heatmaps visualizing the plastically condensed collagen imaged by reflectance confocal microscopy. Scale bar, 200 μm. (F) Dynamics of collagen area change (%) over 48 hours and after the decellularizing process. Data points: mean±s.e.m. (G) The intensity of collagen reflectance imaging from the same batch of collagen gels was measured. Data points: mean±s.d. For (F, G), N = 4 gels per condition from one experiment. (H) Total compaction (elastic+plastic, before decellularization) and plastic compaction (after decellularization) of collagen under varied compression measured by collagen area change. N = 6, 4, 5, 4, and 8 gels per condition from one (for 1% PEG and 3% PEG) or three independent experiments (for Isotonic, 2% PEG, and 4% PEG). Data points: mean±s.e.m. For (G) and (H), the shown statistical comparisons were made between the “Isotonic” condition and the other conditions. Statistic significance was calculated by one-way ANOVA with a Tukey multiple comparisons test (*p<0.05; ****p<0.0001; ns: not significant).

Collagen compaction assays are often used to evaluate cell forces in a soft ECM context. Here, we find volumetric compression also influences the dynamics of collagen gel compaction (Fig. 3E, F). At all levels of compression, the collagen gel area reached the plateau in less than 24 hours. With the increase in compression, the level of collagen compaction decreases. Given that collagen is a highly viscoplastic material, we evaluate the plastic deformation of the gels by completely removing cellular force with the addition of 2% Triton X-100 at Hour 48. After the relaxation of elastic deformations caused by cell contraction, we show that majority of the collagen compaction is plastic (Fig. 3F, H). When exposed to the compression, the plastic compaction of collagen (Fig. 3H) and the local collagen fiber condensation (Fig. 3E, third row, and Fig. 3G) is reduced, as a function of the PEG concentration. Comparing isotonic and 2% PEG conditions, we find that although cells are able to reach the same level of actomyosin contractility based on traction forces (Fig. 3C), the compressed cells are not able to compact the gel at the same level. This implies that non-elastic ECM remodeling capabilities are altered by compression and that standard TFM measurements do not reveal these features.

Cellular force plastically remodels collagen ECM^45,46^. We have previously shown dynamic protrusions of cells are required for continuous recruitment and plastic densification of collagen^6,47^. Cells build up both tension and adhesion ligands in fibrous ECM and stiffen the ECM via force and protrusions^48–50^. Considering volumetric compression shifts Rac/Rho balance and controls cell morphodynamics (Fig. 2), we hypothesize the shifted Rac/Rho-regulated cytoskeleton dynamics alter cell-ECM interactions and matrix remodeling under compression. We proceed to investigate the cell-substrate interactions at the single-cell level (Fig. 4). First, we compare the cell spreading state on fibrous collagen gels (gel thickness: ~200μm) and elastic polyacrylamide (PA) substrates (Fig. 4A,B) after seeding under compression for 24 hours. Consistent with previous findings^28,51^, on elastic substrates cell spreading and YAP nucleus translocation are stiffness-dependent (Fig. 4, Supplementary Fig. 4D). On the soft gel (shear modulus ~5.5 Pa, at 1Hz, Supplementary Fig. 3), the cells acquire minimal area and stay rounded (high circularity) with or without the volumetric compression (Fig. 4A,B and Supplementary Fig. 4 A-C). On the stiff gel (shear modulus ~ 7.5 kPa, Supplementary Fig. 3), the cells spread but the spreading area reduced when compression increased. From high-resolution images, we notice the round, non-spreading cells on a soft elastic substrate in the isotonic condition still generate small VASP-rich protrusions while the compressed cells cannot, possibly due to the suppressed Rac activity (Supplementary Fig. 4C). When seeded on a non-elastic collagen surface (shear modulus ~33 Pa, Supplementary Fig. 3) with an initial stiffness of the same order of magnitude as the soft elastic PA gels’ stiffness, the cells are able to spread (Fig. 4A,B) and strongly engage with the substrate by strain-stiffening and continuous plastic collagen fiber recruitment in the isotonic condition (Fig. 4e,f, Supplementary Video 3). Under volumetric compression, however, the cells remain round and cannot drive cell spreading on the collagen surface (Fig. 4A,B,G,H). However, when we crosslink collagen via glycation, we partially recover cell spreading under 4% PEG compression (Fig. 4C,D), presumably via increasing substrate stiffness and reducing force-induced relaxation^47^. The results altogether suggest that volumetric compression does not impair the stiffness- and tension-sensitive actomyosin contractile machinery but rather suppresses the dynamic cytoskeletal protrusion-mediated processes, which are important in the local densification and stiffening of soft non-elastic ECM substrates and the subsequent spreading of cells on the mechanically remodeled environments (Fig. 4E-H). Furthermore, when we remove the contractility of isolated single cells on collagen by depolymerizing actin with Cytochalasin D (CytoD) after cell spreading (Hour 24), the collagen elastically recoiled, leaving behind different levels of plastic collagen densification (Fig. 4I). In the isotonic case, the cells introduce a significant amount of long-range densification of collagen as well as a high strain (total deformation/cell area) due to continuous fiber recruitment and cell spreading, while the highly compressed (4% PEG) cells generate a much shorter range of plastically densified fibers with very limited elastic deformation and strain on the ECM (Fig. 4I,J). Interestingly, the elastic collagen recoil, as well as the strain, in the 2% PEG condition is on par with that in the isotonic condition (Fig. 4J). This implies that the 2% PEG compressed cells still generate a considerable amount of contraction despite a reduction in plastic remodeling via reduced dynamic protrusion.

**Figure 4.**
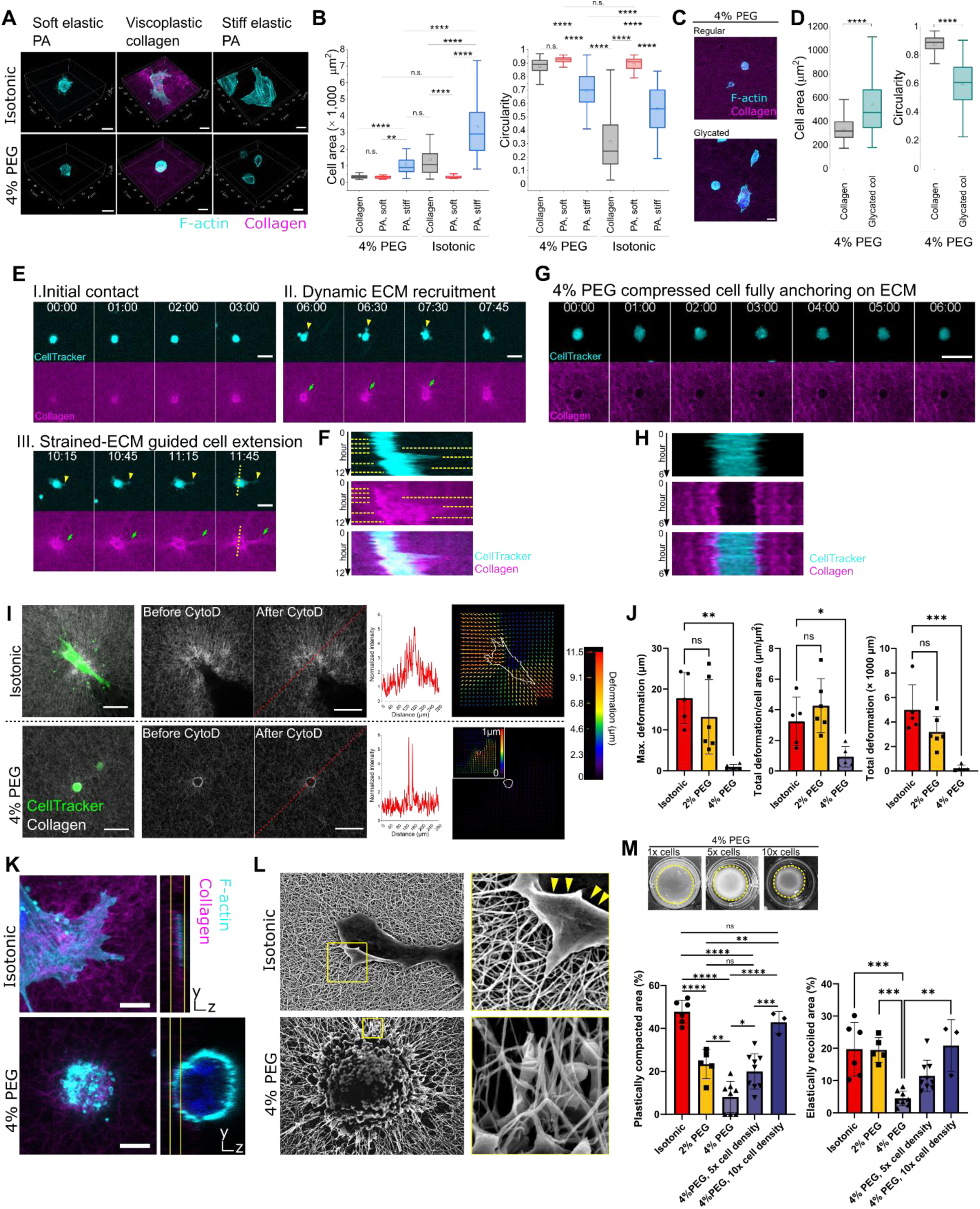
Volumetric compression dictates tension equilibrium of the cell-nonelastic fibrous substrate system via cytoskeleton dynamics. (A) 3D-rendered fluorescent images of non-compressed vs compressed cells seeded on a soft PA gel, a collagen substrate, and a stiff PA gel with the cell area and cell circularity measurement shown in (B). (C) Fluorescent images of 4% PEG compressed cells seeded on the non-crosslinked collagen substrate vs the glycated collagen substrate. The cell area and circularity measurements are shown in (D). Scale bars for (A) and (C), 20 μm. (E) Time-lapse imaging (12 hours, Supplementary Video 3) shows three stages of cell spreading on a collagen substrate: (I) Initial contact: short-range collagen remodeling enables cell anchorage; (II) Dynamic ECM recruitment: the fully anchored cell generates longer protrusion to recruit collagen in the further range. (III) Strain-induced long-range ECM remodeling: continuous collagen recruitment aligns collagen into stiff tracks to guide polarized cell spreading. (F) Kymograph showing cell-ECM interactions from the same 12-hour time-lapse imaging. Yellow dotted lines indicate the protrusions generated from the cell. (G) Time-lapse imaging (6 hours, Supplementary Video 4) of a 4% PEG compressed cell interacting with the collagen substrate, and the kymograph is shown in (H). Scale bars in (E) and (G), 50 μm. (I) Comparing the remodeling and deformation in collagen for 24 hours by cells in isotonic or 4% PEG conditions, before and after removing cell contractility with cytochalasin D (CytoD, 25 μM). The red dotted lines profile collagen density after the CytoD treatment. The displacement fields of collagen were calculated by comparing the collagen substrates before and after the CytoD treatment. The inset shows the collagen displacement field under a different scaling. Scale bars, 100 μm. (J) Elastic recoil of collagen after the cell force removal was characterized by maximum collagen deformation, collagen deformation per unit cell spreading area, and total collagen deformation. N= 5, 6, and 5 cells per condition from one experiment. (K) High-resolution confocal imaging shows local protrusion-ECM interactions and the side profile of the cells on collagen under isotonic vs compressive conditions. Scale bars, 10 μm. (L) SEM images detailing the cell protrusion types and the cell-ECM interactions. Yellow arrowheads indicate membrane ruffling. (M) Collagen compaction assay characterizing plastic ECM remodeling and normalized elastic recoiling under 4% PEG compression at different cell densities (1× cells: 150,000 cells/mL; 5× cells: 750,000 cells/mL; 10× cells: 1,500,000 cells/mL) compared with isotonic and 2% PEG condition at the 1× cell density. N=6, 5, 8, 9, and 3 gels per condition from 3, 2, 4, 4, and 2 independent experiments, respectively.

High-resolution confocal imaging (Fig. 4K) and scanning electron microscopy (Fig. 4L) further confirm that cells make use of dynamic, far-reaching protrusions to spread on a soft, fibrous collagen surface, and reveal that the compressed cells only form short-range, bleb-like protrusions to interact with the local fibers and reach the spreading equilibrium once the local fibers are plastically recruited (Fig. 4G,H; Supplementary Video 4). When shortening the cell-cell distance by increasing cell seeding density (5- or 10-fold increase) in the collagen compaction assay (Fig. 4M), we demonstrate that a larger collection of the short-range plastic recruitment of the 4% PEG compressed cells is still sufficient to achieve large-scaled plastic compaction of the whole gel. A tenfold increase in cell numbers under the 4% PEG compression leads to a comparable compacted gel area to the gels in the isotonic case. This confirms the importance and impact of cell protrusion dynamics in non-elastic cell-ECM interactions.

### Forcing the activation of Rac1 restores compressed cells’ spreading, ECM recruitment via dynamic protrusions, and YAP nuclear translocation

The feedback circuit between cell protrusion, cell spreading, and long-range ECM remodeling (Fig. 4E,F) is impaired by volumetric compression through a shifted Rho/Rac balance in a contractile cell. On 2D rigid surfaces, the cells undergo a transition from “dynamic” to “constrained” due to inhibited Rac1 and enhanced RhoA activity (Fig. 1,2, Supplementary Fig. 1). Here, we demonstrate the capability of partially reversing compressed cell phenotypes by overriding the compression-induced Rho/Rac circuitry. We apply two strategies (Fig. 5A): (1) relaxing Rho/ROCK induced actomyosin contractility via the small molecule ROCK inhibitor (Y27632); or (2) enhancing Rac1 activity via constitutively active Rac (RacV12) or a small GTPases activator. We first investigate cell spreading on the rigid surface. With the inhibition of RhoA’s downstream effector ROCK, we recover cell protrusions and spreading of the compressed cells (Supplementary Fig. 5, Supplementary Video 5). On the Rac1 axis, we find that cells positively transfected with constitutively active Rac (RacV12) exhibit increased cell spreading with edge ruffling under 4% PEG compression (Fig. 5B). Treating 4% PEG compressed cells with a small GTPases activator CN04 also recover lamellipodia formation and protrusion dynamics (Fig. 5C,D, Supplementary Video 6), as well as cell spreading and cell shape (Fig. 5E). We confirm that CN04 induces long-lasting (at least 6 hours) effects on Rac1 activation but not on RhoA (Supplementary Fig. 6). To further rule out the possibility that potential activation of Rho by CN04 contributes to cell spreading, a Rho activator (CN03) is used. We show that solely activating RhoA fails to induce changes in actin organization, cell area, and cell shape in the compressed cells (Fig. 5C). Interestingly, although the GTPase activator CN04 has prolonged activation on Rac1, the reactivated protrusion dynamics seem to reduce within 2 hours but the cell continues to spread and eventually reaches the steady state at a higher cell area (Fig. 5D,E). This may suggest a biphasic effect of Rac on protrusion dynamics; Rac1 is necessary for protrusion generation but increasingly activated Rac1 over time can also counteract Rho activity^34^, which impedes the retraction of cell protrusions.

**Figure 5.**
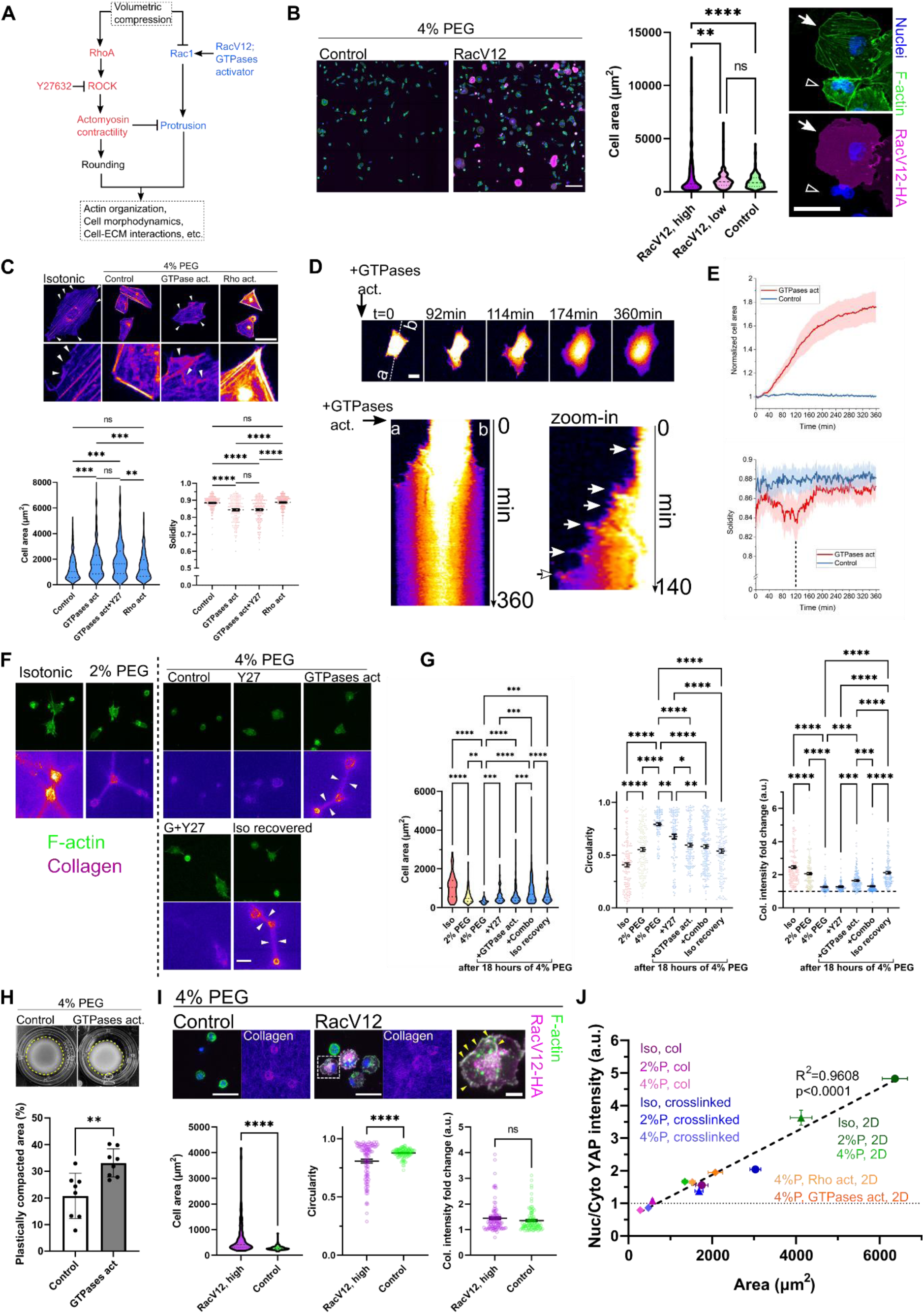
Restoring Rac activity in highly compressed cells. (A) Schematic indicating the hypothesized approaches to rescue the cell behaviors altered by compression. (B) Fluorescent imaging of the cells transfected with constitutively active Rac (RacV12) and the control counterpart. Cell area was measured to evaluate cell spreading. The close-up images on the right panel show two cells, of which only one was transfected with RacV12 (indicated by the arrow) and showed enhanced spreading and ruffling edges. The non-transfected counterpart (indicated by the hollow arrowhead) exhibits distinct peripheral actin bundles. Scale bars, 200 μm (left panel) and 50 μm (the close-up images, right panel). (C) Actin organization (shown as the intensity heatmap), cell spreading area, and cell solidity on the 2D rigid surface under 4% PEG compression with vehicle (Control) and three drug treatments: GTPases activator (GTPases act, CN04), a combination of GTPases activator and ROCK inhibitor Y27632 (Y27, 1μM), and Rho activator (Rho act, CN03). N=155, 142, 179, 216 cells for the corresponding conditions from two independent experiments. Mean±s.e.m. (D) Time-lapse imaging showing the morphodynamics of 4% PEG compressed cells upon GTPases activation (Supplementary Video 6). Scale bar, 20 μm. Temporal measurements of cell area and solidity of control cells and the activated cells are shown in (E). For (E), n=9 and 12 cells for Control and GTPases activation respectively. (F) Imaging of F-actin of cells spreading on fluorescently-labeled collagen cushion under varied compression conditions and drug treatments. “G+Y27” indicates the combined treatments of GTPases activator+Y27632 (1μM). Scale bar, 50 μm. Cell area, cell circularity, and collagen densification underneath the cells (intensity fold change compared to the background) were measured for the corresponding conditions in (G). N=143, 160, 90, 115, 145, 193, and 140 cells for the corresponding conditions from two to three independent experiments. Mean±s.e.m. (H) Collagen contraction assay (cells density: 750,000 cells/mL) characterizing plastic ECM remodeling under 4% PEG compression with and without the GTPases activation. Mean±s.d. (I) Cells transfected with constitutively active Rac (RacV12) and the control counterparts seeded on collagen cushions. The close-up image shows a RacV12 positive cell and its edge ruffling indicated by the arrowheads. Cell area, cell circularity, and collagen densification under the cells were measured. N=105 RacV12-transfected cells, and 97 control cells from two independent collagen cushions. Mean±s.e.m. Scale bars, 50 μm; scale bar in the close-up inset, 10 μm. (J) Correlation between cell spreading area and YAP nuclear translocation cross compression conditions (Iso: isotonic; 2%P: 2% PEG; 4%P: 4% PEG), drug treatments, and substrate types (2D: collagen coated rigid surface; col: collagen cushion; crosslinked: glycated collagen cushion). Statistic significance was calculated by one-way ANOVA with a Tukey multiple comparisons test for (C) and (G). Student t-test was used in (H) and (I). (*p<0.05; **p<0.01; ***p<0.001; ****p<0.0001; ns: not significant) In (J), statistical differences between conditions are shown in Extended Data Fig. 7.

**Figure 6.**
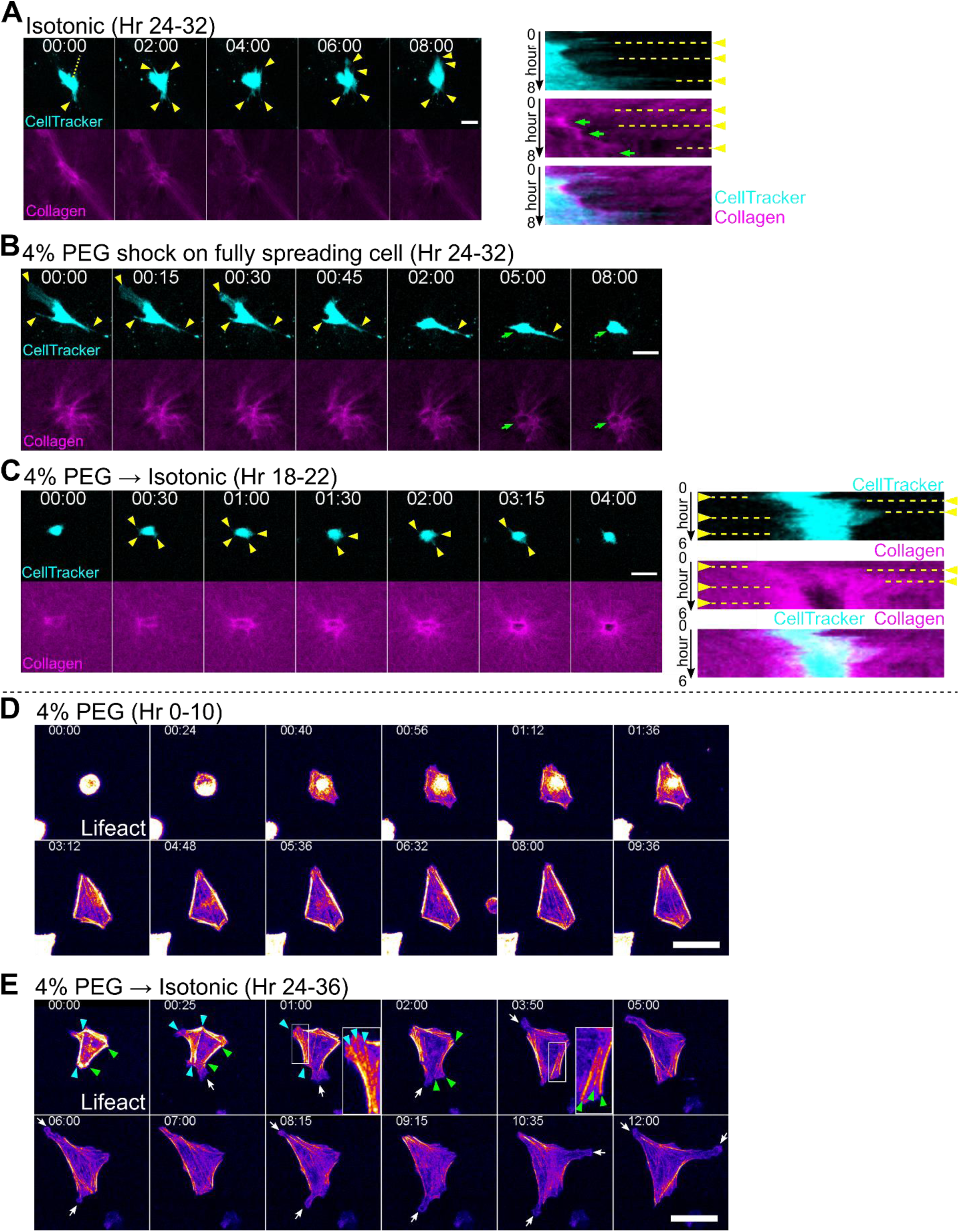
Directing cell cytoskeletal states reversibly via volumetric compression. (A) Time-lapse imaging (8 hours, Supplementary Video 8) of a fully anchored cell interacting with the surrounding collagen via the dynamic protrusions (marked by the arrowheads). The kymograph along the dotted line shows the extension/retraction of the protrusions and the gradually densified collagen (green arrows) by the protrusions over time. (B) Volume compression shock dewets a fully spreading cell on collagen (Supplementary Video 9). Cell protrusions are marked by yellow arrowheads. The green arrow indicates the contractile cell cortex anchoring on the collagen cushion. (C) Time-lapse imaging of a compressed cell anchored on collagen recovering for 8 hours in the isotonic condition (Supplementary Video 10). Recovered cell protrusions (yellow arrowheads) rapidly recruit and densify the surrounding collagen fibers. Kymograph shows the fluctuation of the cell boundary and protrusions while accumulating collagen around the cell. (D) Actin organization of a 4% PEG compressed cell spreading on a collagen-coated rigid surface. Pronounced peripheral actin bundles emerged from the cell actin cortex. (Supplementary Video 11) (E) A compressed cell on a rigid surface recovering in the isotonic condition shows redistribution of stress fibers from two sets of peripheral actin bundles (indicated by arrowheads in cyan and green, respectively) and the freed dynamic protrusions (white arrows) from the anchorage points. (Supplementary Video 13) Scale bars in (A-E), 50 μm.

On soft fibrous ECM, we show that reshaping the Rac/Rho balance in the compressed cells by small molecule drugs also switches cell spreading and cell-ECM interaction phenotypes (Fig. 5f,g, Supplementary Fig. 7). Both Rac1 activation (CN04) (Supplementary Video 7) and ROCK inhibition even at low dosing (Y27632, 1μM), but not Rho activation, can rescue cell spreading in the compressed cells. Notably, the rescued cell spreading by relaxing cell contractility can be inhibited by the Rac1 inhibitor NSC23766 (Supplementary Fig. 7), suggesting an inhibitory role of Rho/ROCK-regulated contractility in Rac1-directed protrusion formation. By measuring fold change in collagen intensity underneath the cells, we find that only Rac1 activation recovers ECM densification and long-range remodeling (Fig. 5F,G). The co-treatment of Rac1 activation by CN04 and ROCK inhibition leads to a more drastic recovery of cell spreading but collagen densification is diminished. At the bulk level, we show that activating Rac1 with the GTPase activator in the 4% PEG compressed cells promote collagen plastic compaction (Fig. 5H). Transfecting the cells with constitutively active Rac also rescues cell spreading on the soft fibrous ECM under 4% PEG compression but fails to increase collagen densification (Fig. 5I). Although thin edge ruffling is observed in the RacV12-transfected cells, the lamellipodia formation via the forced Rac activation is likely driven by the actin growth without modulating cell contraction^52^. This suggests that Rho/ROCK-mediated cell contractility and cell protrusion-retraction dynamics are necessary for establishing ECM tension and dynamic fiber recruitment, but ROCK inhibition (Fig. 5F,G) or solely activating Rac (Fig. 5I) are sufficient to recover tension-independent cell spreading in highly compressed conditions. Furthermore, by comparing YAP nuclear translocation across some conditions where cells can recover spread area while maintaining contractility, including Rac1 activation and collagen cross-linking, we show that compression regulates YAP activity via cell spreading area (Fig. 5J and Supplementary Fig. 8). By recovering cell spreading via matrix stiffening (crosslinking) or activating Rac1 without disrupting cell contractility, we demonstrate a partial rescue of YAP nuclear translocation under high compression. Notably, solely activating RhoA with a suppressed cell spreading under compression is not sufficient to trigger YAP nuclear translocation.

**Figure 7.**
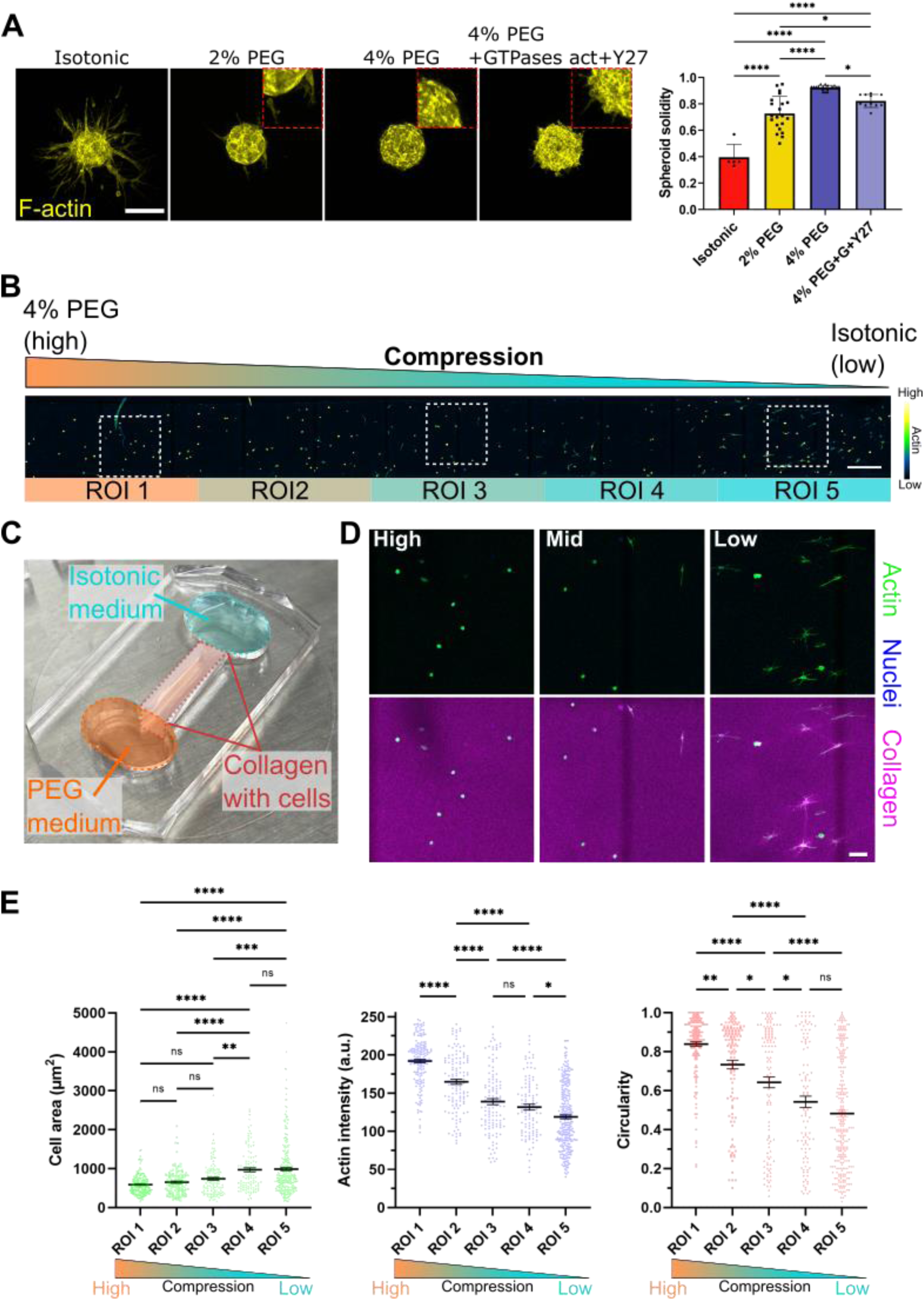
Volumetric compression regulates cell-ECM interactions in 3D. (A) Staining of F-actin after 48-hour collagen invasion of multicellular spheroids in conditions of isotonic, 2% PEG, 4% PEG, and 4% PEG with the “GTPases activator+Y27” treatment. The solidity of the spheroids was used to evaluate the fluidity of the cells. Scale bar, 200 μm. n=5, 21, 22, 11 spheroids for the corresponding conditions, from three to four independent experiments. Statistic significance was calculated by one-way ANOVA with a Tukey multiple comparisons test. (*p<0.05; **p<0.01; ***p<0.001; ****p<0.0001; ns: not significant) (B) We introduce a continuous volumetric compression gradient by a PEG concentration gradient throughout a collagen gel with SNU475 cells with a device depicted in (C). Scale bar in (B), 500 μm. The diameter of the cover glass used for the device is 40 mm. (D) Representative images of cells embedded in collagen gel, experiencing varying levels of compression (Max. projection of 410 μm thickness). Compression suppresses the generation of cell protrusions and thus disenables long-range ECM remodeling. Scale bar, 100 μm. (E) We divide the osmotic gradient into five regions of interest (ROI): ROI 1-5. We quantify cell area, actin intensity, and circularity in each region. These three features of the cell exhibit a gradient, in response to the introduced osmotic gradient. Overall, we demonstrate the tunability of cell dynamics and non-elastic cell-ECM by controlling the environmental osmolarity. n=172, 123, 122, 106, and 276 cells for the ROI1 to ROI5 respectively, pooled from two independent devices. Statistic significance was calculated by one-way ANOVA with a Tukey multiple comparisons test. (*p<0.05; **p<0.01; ***p<0.001; ****p<0.0001; ns: not significant)

**Figure 8.**
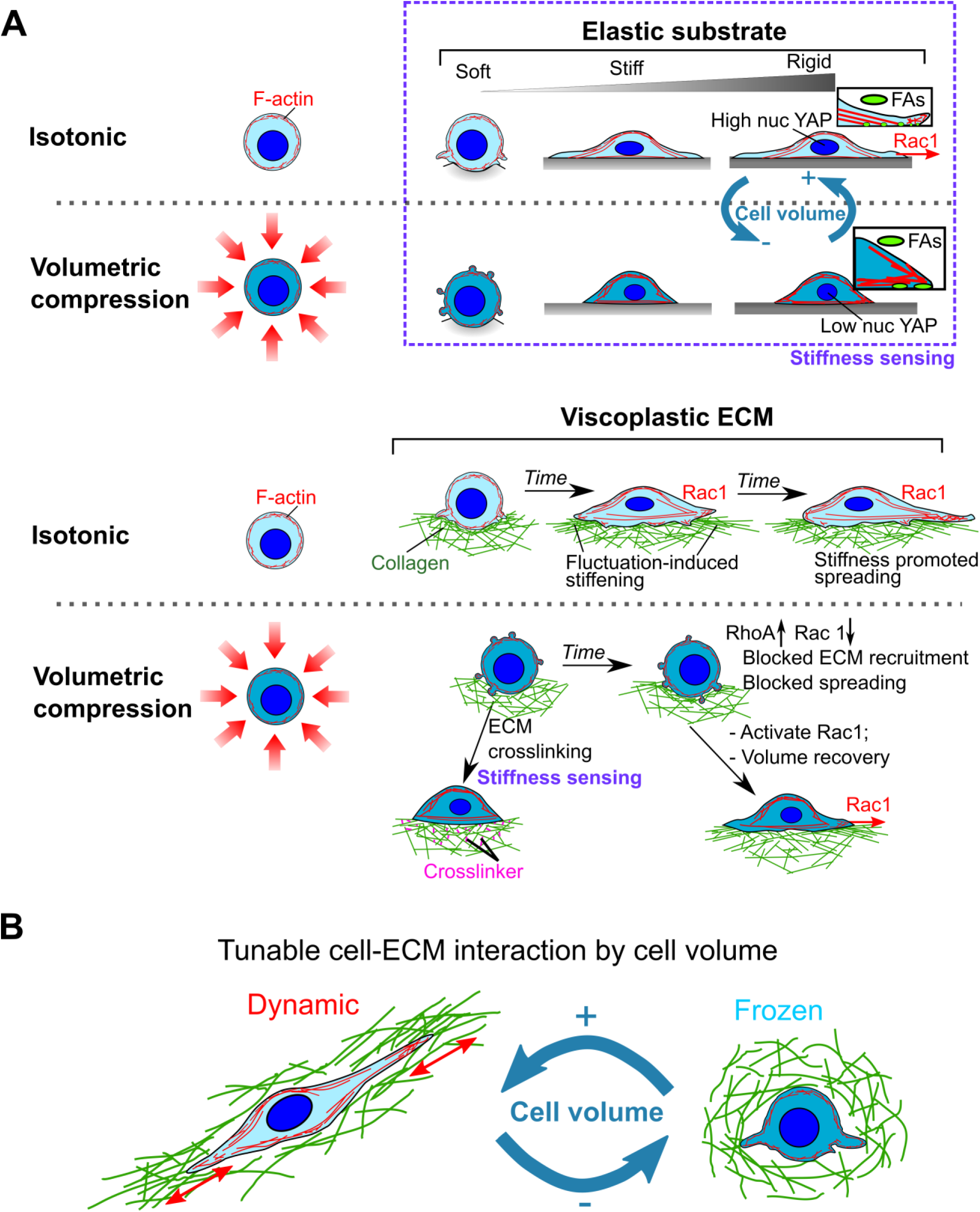
Proposed model of the mechanosensing and cell-ECM nonelastic interactions dictated by Rho/Rac rebalancing via volumetric compression. (A) Shifted cell mechanosensing under volumetric compression. The intact actomyosin machinery enables cells to spread via stiffness sensing on elastic, rigid surfaces with a changed arrangement of cytoskeleton and FA with a conserved total FA area. However, on the soft non-elastic collagen the suppressed dynamic protrusions fail to initiate the long-range fiber recruitment thus the strain stiffening in the ECM to facilitate further cell spreading, which can be reverted by re-activating Rac1 in the compressed cells. Due to the intact actomyosin machinery and rigidity sensing, increasing ECM tension by collagen fiber crosslinking is also sufficient to partially rescue the suppressed cell spreading under compression. Furthermore, this volumetrically controlled cell mechanical state by osmolarity is reversible by cell volume recovery. (B) Controlling cell volume programs cell-ECM nonelastic interactions in 3D via protrusion dynamics.

### Rapid switching between actin cytoskeletal states can be controlled by volumetric compression

Consistent with a previous study^53^, the cells allowed to spread for 24 hours in the isotonic condition adopt a polarized morphology with high ECM remodeling in their vicinity. Interestingly, the morphodynamics and protrusions continue enabling ECM recruitment in the less remodeled region over 8 hours (Fig. 6A, Supplementary Video 8). When shocking a fully spreading cell on collagen with 4% PEG (Fig. 6B, Supplementary Video 9), we observe that the cell’s morphodynamics is almost instantly suppressed, leading to a gradual retraction of the established long-reaching protrusions, and eventual cell rounding with a local “condensed ring” of the collagen underneath. Having demonstrated that recovering the highly compressed cells in isotonic media restores cell spreading and ECM recruitment (Fig. 5F,G), time-lapse imaging further confirms that the compressed cells release dynamic protrusions to condense and align the surrounding ECM fibers when reverted back to isotonic media (Fig. 6C, Supplementary Video 10). Together, the results suggest that the volumetric compression effects on morphodynamics and cell protrusions are reversible.

We then investigate the implications of this reversible effect on the actin cytoskeleton using time-lapse imaging on 2D rigid surfaces (Fig. 6D,E). We utilize Lifeact-RFP expressing SNU475 cells to label actin. These cells exhibit consistent temporal spreading dynamics as observed in the Celltracker labeled cells (Fig. 1 and Fig. 2) under 4% PEG compression. When spreading in the 4% PEG medium, a gradual transition from the actin cortex to peripheral actin bundles can be seen during cell spreading (Fig. 6D, Supplementary Video 11). With minimum lamellipodia observed, the extension of these thick actin bundles mainly drives the cell spreading, possibly via adhesion. After 4-5 hours, the cell area and morphology reach a steady state with a stable and almost “frozen” actin organization, which resembles the early stage of cell spreading in isotonic conditions (“Initial contact” stage, Supplementary Fig. 9 and Supplementary Video 12). When released back into isotonic media from 24-hour compression (Fig. 6E, Supplementary Video 13), the compressed cells rapidly deploy protrusions, which is also recapitulated by the spreading stage in the isotonic condition (“Spreading” stage, Supplementary Fig. 9). These newly developed protrusions evolve into sheet-like lamellipodia and facilitate the splitting of the existing thick peripheral actin bundles into multiple thinner actin cables, which resemble typical transcellular stress fibers. We thus discover an intrinsic relationship between the cell cortex, thick peripheral actin cables, and transcellular stress fibers, which are inter-convertible by dynamic Rac1-driven actin protrusions tuned by cell volume. Notably, the restored dynamic protrusions in these “unfrozen” cells growing typically from the vertices of the peripheral actin bundles suggest an inhibitory effect of membrane-proximal actin on cell protrusion formation^54^. These findings highlight a central role of dynamic protrusions in cytoskeletal architecture patterning and redistribution.

Alongside others^19,55^, using multicellular tumor spheroids (MCTS), we demonstrate that fully encapsulated cells in a collagen matrix (3D) capture the same phenotype under compression as seen on the collagen cushions (Fig. 7A). Volumetric compression sufficiently induces mesenchymal cells to transform into a non-motile state and block the “solid to fluid” phase transition^56^, characterized by high solidity of the spheroid shape. By rebalancing Rho/Rac activities via Rac1 activation and ROCK inhibition, we partially recover invading protrusions, measured by a reduction in the spheroid solidity. We also develop a cell culture chip that introduces a continuous osmotic gradient to single cells embedded in 3D collagen (Fig. 7 B,C). We find actin arrangement, actin intensity, and collagen remodeling are functions of the level of environmental osmolality (Fig. 7 D,E), further demonstrating that cell mechanical states can be tightly tuned by cell volume. To demonstrate the universality of the volumetrically controlled cytoskeletal states and YAP translocation cross multiple dynamic cell types, we also volumetrically compress mouse mesenchymal stem cells (mMSCs) (Supplementary Fig. 10A-D), metastatic breast cancer cells MDA-MB-231 (Supplementary Fig. 10E-H), and human umbilical vein endothelial cells (HUVECs) (Supplementary Fig. 11). After 24 hours of spreading on the rigid surface under compression, all the cell types exhibit polygonal shapes consisting of pronounced peripheral actin bundles and smaller cell area. Interestingly, when losing far-reaching dynamic protrusions under compression, the connectivity of the endothelial cell network is reduced in the tube formation assay on Matrigel (Supplementary Fig 11E,F). By relaxing cell contractility with a ROCK inhibitor, we are able to recover the suppressed protrusions and re-establish the endothelial network.

## Discussions and conclusion

In this study, we demonstrate that volumetric compression is a biophysical master regulator of mechanobiological cell states and functions, notably cytoskeletal organization and mechanical performance, elastic and non-elastic cell-matrix interactions, and mechanosensing (Fig. 8). The underlying mechanochemical circuitry is centered on tuning the balance of Rho GTPase activities, highlighting the programmability of these signals by volume regulation.

Actin organization and structures play functional, mechanical roles. We demonstrate that volumetric compression modulates cytoskeletal states, as well as the distribution of adhesions, reversibly. When compressed, interior stress fibers are significantly reduced and peripheral/cortical actin cables become more prominent, coupled with a loss of lamellipodia and cell boundary fluctuations. In 2D, this is associated with altered adhesions from smaller, evenly distributed plaques to bigger, more clustered adhesion sites that are primarily localized at the cell periphery. Despite the redistribution of cell adhesion, the total adhesion area is conserved. Thus, volumetric compression renders cells to be more polygonal, compact shaped with minimum motility, which we term as a “frozen state”. Beside interrogating the volumetric compression, we also report a similar peripheral actin bundle arrangement in response to the mechanical compression applied onto the cells (Supplementary Fig. 12), suggesting a universal cellular response to compressive stresses. Architectural changes in actin can be triggered in response to environmental changes, such as cell confluency^30^, to maintain cohesion and tension across epithelial sheets^57^. Our study isolates the effect of cell volume on cytoskeletal states in single cells and reveals how actomyosin and adhesion machinery respond to volumetric compression. Recent work demonstrates that actin stress fibers can be embedded within the cell actin cortex, and these two actin architectures share synergistic actions in cell contractility ^58^. In our study, we show that interior stress fibers and thick peripheral cables can originate from the same actin bundles that can be interconverted via volumetric compression. Specifically, volumetric compression modulates lamellipodia and cell boundary fluctuation dynamics, which mediate the redistribution of stress fibers from within thick peripheral cables.

In ECMs, cell protrusions can play mechanical roles in cell migration and spreading by non-elastically remodeling the local matrix, e.g. mechanically opening pores or locally accumulating matrix fibers^6,59^. Our study demonstrates that volumetric compression regulates cytoskeletal organization and fundamentally alters its mechanical function. Notably, overall cell contractility, which is a common measure of cell mechanical output, does not fully recapitulate the altered states and function of compressed cells, especially in the context of cell-matrix interactions and the cell-induced compaction of non-elastic gels. While cell traction forces are comparable between compressed cells and non-compressed cells (especially in 2% PEG vs. isotonic), ECM compaction at the global scale and densification at the local scale are significantly reduced under compression (Fig. 3). These non-elastic interactions are associated with the dynamic fluctuations of the cell periphery, highlighting the importance of cells dynamically “sampling” their surroundings in a complex mechanical environment. These findings provide a potential approach to pattern non-elastic multicellular systems (e.g. *in vitro* microtissues^60^) by tuning cell-ECM interactions via modulating osmolarity (e.g. a continuous osmotic gradient in 3D matrices, Fig. 7B-E).

The interplay of cells and their substrates entails not only spontaneous mechanical interactions but also feedback through mechanosensing. Recent findings showed the critical impact of substrate stiffness^61,62^ and Rac1-mediated rigidity sensing^62^ for tumorigenic cell reprogramming. In addition to stiffness, cells can simultaneously exhibit compressed profiles in complex diseased tissues such as expanding tumors (Fig.1B). Our results show that the interplay between compression and substrate properties modulates mechanosensing phenotypes and subsequent cell and tissue states. On soft, non-elastic collagen substrates, cells can mechanically densify their surroundings, which results in increased ligand density and stiffness. Cells can then sense the remodeled environment and respond accordingly via spreading and transcriptional programs. Compressed cells, however, have impaired ability to mechanically remodel non-elastic surroundings due to suppressed protrusion dynamics, resulting in reduced cell spreading and YAP nuclear translocation on these substrates (Fig. 5J, and Supplementary Fig. 7). However, compressed cells are still able to respond expectedly to stiff substrates (e.g. crosslinked collagen or stiff PA substrates), indicating that their canonical stiffness-sensing pathways, which are primarily driven by actomyosin contractility and adhesions, are intact (Fig. 4A-D). Our findings highlight the important role of dynamic cell boundary fluctuations, particularly lamellipodial-like dynamics, as the initiator of cell spreading and tension building, as well as the mediator of mechanosensing, in non-elastic substrates. Volumetric compression, therefore, acts as a mechanosensing switch in non-elastic tissue microenvironments.

Our data demonstrate that the impact of volumetric compression on cell states is driven at the molecular signaling level by the rebalancing of Rho GTPase activities, particularly Rac1 and RhoA. Compression suppresses Rac1 and activates RhoA, shifting their equilibrium ratio, leading to the reduction of Rac1-driven cell spreading and protrusion dynamics while maintaining RhoA-driven contractility. This ratio can be subsequently biochemically controlled to override mechanical regulation of cell states and mechanosensing phenotypes. Reverting Rho GTPase activities by Rac1 activation or ROCK inhibition unfreezes compressed cells and partially rescues protrusion-related cellular functions, such as cell spreading, boundary ruffling, invasive protrusions, and endothelial network formation. In summary, our study reveals the mechanisms and consequences of cell state programming via volumetric compression.

## Acknowledgments

We thank Dr. Jing Yan for the access to the rheometer. We also thank Dr. Xuchen Zhang from Yale Surgical Pathology for the histology samples of liver cancer patients. We acknowledge Saltzman Lab for the access to their plate reader. We thank Dr. Martin Schwartz for the RacV12 plasmid. We are also grateful for the valuable discussion with Dr. Michael Murrell on the TFM experiments. We thank Dr. Rong Fan for the access to their EVOS microscope. We are supported by: Yale Liver Center Pilot Project under award NIH P30 DK034989 (M.M.), NIH R35GM142875 (M.M.), NIH grant T32EB019941 (R.Y.N).

## Resource availability

### KEY RESOURCES TABLE

| REAGENT or RESOURCE | SOURCE | IDENTIFIER |
| --- | --- | --- |
| Antibodies |  |  |
| anti-YAP | Santa Cruz Biotechnology | Cat#sc-101199 |
| anti-Myosin light chain (phospho S20) | Abcam | Cat#ab2480 |
| anti-VASP | Santa Cruz Biotechnology | Cat#sc-46668 |
| anti-MYO10 | Abnova | Cat#PAB21665 |
| anti-hemagglutinin (HA) tag-Dylight 488 | Invitrogen | Cat#26183-D488 |
| Alexa Fluor 488 Goat Anti-Mouse IgG antibody | Invitrogen | Cat#A-11001 |
| Alexa Fluor 647 Goat Anti-Rabbit IgG antibody | Invitrogen | Cat#A-21244 |
| Chemicals, Peptides, and Recombinant Proteins |  |  |
| Hoechst 33342 | Invitrogen | Cat#H3570 |
| Rhodamine Phalloidin | Abcam | Cat#ab235138 |
| Alexa Flr 488 Phalloidin | Invitrogen | Cat#A12379 |
| CellTracker Green | Invitrogen | Cat#C7025 |
| Y27632 | Cayman | Cat#10005583 |
| NSC23766 | Tocris | Cat#2161 |
| Cytochalasin D | Abcam | Cat#ab143484 |
| Rho activator CN03 | Cytoskeleton, Inc. | Cat# CN03 |
| Small GTPases (Rho/Rac/Cdc42) activator CN04 | Cytoskeleton, Inc. | Cat# CN04 |
| 3-hydroxytyramine hydrochloride | Acros Organics | Cat#122001000 |
| RPIM 1640 | Corning | Cat#15-040-CV |
| GlutaMAX | Gibco | Cat#35050061 |
| Fetal bovine serum | Gibco | Cat#26140079 |
| Penicillin-Streptomycin | Gibco | Cat#15140122 |
| 0.25% Trypsin-EDTA | Gibco | Cat#25200056 |
| Microvascular Endothelial Cell Growth Medium | Cell Applications | Cat#111-500 |
| Collagen I, High Concentration, Rat Tail, 100 mg | Corning | Cat#354249 |
| Polyethylene glycol 300 | Sigma | Cat#8.07484 |
| Alexa Fluor 647 NHS Ester dye | Invitrogen | Cat#A20006 |
| D(-)-Ribose | Acros Organics | Cat#A0405792 |
| (3-Aminopropyl)trimethoxysilane | Thermo Scientific | Cat#A1128422 |
| Glutaraldehyde | Polysciences | Cat#01909-10 |
| 40% acrylamide | Sigma | Cat#A4058 |
| 2% *N,N'*-Methylenebisacrylamide solution | Sigma | Cat#M1533 |
| Ammonium persulfate | Sigma | Cat#A3678 |
| *N,N,N,N'*-tetramethylethylenediamine | MP Biomedicals | Cat#194019 |
| Sulfo-SANPAH | Thermo Scientific | Cat#22589 |
| Dichlorodimethylsilane | Sigma | Cat#440272 |
| Carboxylate-Modified Microspheres, dark red | Thermo | Cat#F8807 |
| Triton™ X-100 | Sigma | Cat#T8787 |
| Opti-MEM | Gibco | Cat#31985062 |
| Lipofectamine™ 3000 Transfection Reagent | Invitrogen | Cat#L3000001 |
| Pierce™ BCA Protein Assay Kit | Thermo Scientific | Cat#23225 |
| Sequa-brene | Sigma | Cat#S2667 |
| 4% paraformaldehyde in PBS | Santa Cruz Biotechnology | Cat#sc-281692 |
| Osmium tetroxide, 2% solution | Polysciences | Cat#23310 |
| PBS | Gibco | Cat#10010-023 |
| BSA | Sigma | Cat#A9647 |
| Hexamethyldisilazane | Sigma | Cat#379212 |
| PDMS Sylgard™ 184 | Electron Microscopy Sciences | Cat#24236-10 |
| Puromycin Dihydrochloride | Gibco | Cat#A1113803 |
| Critical Commercial Assays |  |  |
| Rac1 G-LISA Activation Assay Kit | Cytoskeleton, Inc. | Cat#BK128 |
| RhoA G-LISA Activation Assay Kit | Cytoskeleton, Inc. | Cat#BK124 |
| Experimental Models: Cell Lines |  |  |
| SNU475 | ATCC | CRL-2236 |
| Mouse bone marrow mesenchymal stem cells | ATCC | CRL-12424 |
| GFP Expressing Human Umbilical Vein Endothelial Cells | Angio-Proteomie | cAP-0001GFP |
| MDA-MB-231 | ATCC | HTB-26 |
| Recombinant DNA |  |  |
| rLV<sup>Ubi</sup>-LifeAct-TagRFP lentiviral vector | ibidi | Cat#60142 |
| RacV12 plasmid | Driscoll et al. <sup>52</sup> | N/A |
| Software and Algorithms |  |  |
| Fiji | Schindelin et al. <sup>64</sup> | RRID:SCR_002285 |
| Imaris | Oxford Instruments | RRID:SCR_007370 |
| Origin 2020b | OriginLab | RRID:SCR_014212 |
| GraphPad Prism 9 | GraphPad | RRID:SCR_002798 |
| Leica LAS-X | Leica | RRID:SCR_013673 |

## Lead contact

Requests for resource and reagents should be directed to Michael Mak.

## Materials availability

The stably transfected cells expressing RFP-LifeAct generated in this study will be available upon request.

## Data and code availability

All data reported in this paper and additional information required to reanalyze the data will be shared by the lead contact upon request.

## Experimental model and subject details

### Cell lines

Mesenchymal liver cancer cell line SNU475 was cultured in RPIM 1640 (Corning) with 1% GlutaMAX (Gibco), 10% fetal bovine serum (FBS, Gibco), and 1% Pen Strep (Gibco). Mouse MSCs and breast cancer cell line MDA-MB-231 were cultured in DMEM (Gibco) with 10% FBS (Gibco) and 1% Pen Strep (Gibco). HUVECs were cultured in the microvascular endothelial cell growth medium (Cell Applications). Cells were passaged with 0.25% Trypsin-EDTA (Gibco) when confluency reached 80-90%. Cell culture medium was changed every 2-3 days.

## Method details

### Cell culture in compression medium

To prepare the isotonic and the hyperosmotic culture media with a controlled nutrient concentration, we added 4% 1xPBS (isotonic), 2% PEG-300 plus 2% 1xPBS (mid-level compression), and 4% PEG-300 (high-level compression) to the supplemented culture media by volume. All the experiments based on the 2D rigid surface were conducted on collagen I coated plastic surface of 15-well μ-slides (ibidi). The surface was incubated with 50 μg/mL collagen I (High concentration rat tail collagen, CORNING) solution in 0.02N acidic acid at 37°C for 1 hour, followed by washing three times in PBS and air drying. Trypsinized cells were subsequently resuspended in the isotonic or hyperosmotic media and evenly seeded on the rigid, elastic, or collagen substrates.

### Collagen cushion fabrication

For collagen anchorage, the glass surfaces of the 35mm glass-bottom dishes or glass-bottom well plates (Cellvis) were coated with 1.8 mg/mL 3-hydroxytyramine hydrochloride (or dopamine-HCl, Acros Organics) in 10mM Tris-HCl buffer (pH 8.5) for 1.5 hours at room temperature (RT), followed by a thorough wash in cell culture grade water. The coated dishes were then air-dried in a 60 °C oven and cooled down to the RT before use. Round glass coverslips (8mm in diameter) were treated in sterilized 3% bovine serum albumin solution in PBS for 2-3 hours at RT or 4°C overnight, rinsed with water, and air-dried before use. High concentration rat tail collagen (~10mg/mL, CORNING) was neutralized by 0.5M NaOH sterile solution and then adjusted to the final concentration with water and 10x PBS. In this study, the final concentration of collagen is 2mg/mL. A 25-μL neutralized collagen droplet was gently dispensed onto the dopamine-HCl coated glass surface to form a dome without any bubbles. A BSA-treated coverslip was quickly assembled on top of the collagen dome. Due to the surface tension of the collagen solution, the droplet flattened out to cover the coverslip by maintaining a thickness of a few hundred micrometers. After full gelation at 37°C in a humid incubator for 1 hour, the collagen cushions were soaked in PBS for 15-30 min and the top coverslips were then dislodged by gentle PBS streams repeatedly pipetted at the collagen-coverslip interface. To visualize collagen fibers and quantify collagen remodeling, the collagen cushions were incubated in Alexa Fluor 647 NHS Ester dye (12-20 μg/mL) (Thermo) in PBS at 37°C for 1 hour and were thoroughly washed with sterile PBS for three to five times until the buffer was colorless.

### Post-glycation of collagen cushions

We prepared a ribose (Sigma) solution at a concentration of 250mM in PBS. The ribose solution was sterilized with a 0.22 μm syringe filter. Collagen cushions were soaked in 2mL ribose solution per dish at 37 °C in a humid incubator for five days. Before seeding cells on the cushion, the collagen cushions were thoroughly washed with fresh PBS three times followed by quenching in the PEG-conditioned media.

### Fabrication of polyacrylamide (PA) substrates

To fabricate the soft and stiff PA substrates for cell spreading experiments, we followed the protocol described previously^41^. In brief, clean glass surfaces of 35-mm glass-bottom dishes were coated with a thin, even layer of 0.1N NaOH and then silanized with (3-aminopropyl)trimethoxysilane (Thermo Scientific) for 3 min. After a thorough wash, the glass surfaces were treated with 0.5% glutaraldehyde (Polysciences, Inc.) for 30 min. The dishes were then rinsed with deionized water and dried in a 60 °C oven. A gel premix was made of 250 μL 40% acrylamide (AA) (Sigma) and 125 μL 2% bis-acrylamide (bis-AA) (Sigma). The soft gel is made by thoroughly mixing 20 μL premix and 230 μL PBS, and the stiff gel is made by mixing 75 μL premix and 175 μL PBS. Polymerization of the PA gels was initiated by introducing 0.75 μL N,N,N′, N′-tetramethylethylenediamine (TEMED) (MP Biomedicals, LLC) and 2.5 μL 10% ammonium persulfate (APS) (Sigma). 15 μL of the final mix was dispensed on the silanized glass surface and rapidly covered by a dichlorodymethyl silane (Sigma)-treated square coverslip (12mm×12mm). The gel mix was allowed to polymerize for 30 min before the coverslip was carefully removed. After a 30 min UV sterilizing cycle in sterile PBS, the PA substrates were functionalized with 0.2mg/mL Sulfo-SANPAH (Thermo Fisher) followed by incubation in 100 μg/mL collagen I solution in 0.2N acidic acid at 4°C overnight. The substrates were washed three times with PBS before seeding cells.

### Traction force microscopy

The elastic PA substrates for TFM were prepared and functionalized by the protocol described in the previous sections, except that the gel mix was made followed by the reported recipe^63^: 94 μL 40% AA, 15μL 2% bis-AA, 364 μL PBS, 3.4 μL 0.2-μm dark red carboxylated microbeads (Invitrogen), 5 μL 10% APS and 1 μL TEMED. The shear modulus of the gel was characterized by a rheometer as 3 kPa. SNU475 cells were seeded on the PA substrates at the density of 2000-4000 cells/dish in the isotonic or PEG-conditioned media for 24 hours. Before the experiments, the cells were labeled with CellTracker Green (Invitrogen) for 25 min and washed with the same conditioned medium. Randomly selected isolated single cells and the microbeads substrate were imaged with a 20× objective on a position-saving confocal microscope (Leica SP8). The cells were then removed by carefully adding PBS-diluted Triton X-100 (Sigma) into the dish at a final concentration of 8%. The recorded positions of the microbeads substrates were imaged again after 15 min. The drift between microbeads fields before and after the cell removal was corrected with the plugins “Correct 3D drift” or “Descriptor-based registration” on ImageJ. The microbeads displacement field was calculated by the plugin “PIV” and cell traction force was then calculated and plotted by the plugin “FTTC” using a Poisson ratio of 0.5 and a regulation factor of 3×10^-9^. Strain energy of a single cell was derived using the calculated traction vectors (***T***) and deformation vectors (***u***) by the following definition (Eq. 1):

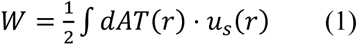

### RhoA and Rac1 activation assays

Colorimetric-based GLISA assays (Cytoskeleton, Inc.) were used to quantify RhoA and Rac1 activity. SNU475 cells (500,000 cells/dish) were seeded on collagen I-coated 100 mm plastic Petri dishes (Falcon) in the isotonic or PEG-conditioned medium. Cells were allowed to spread for 24 hours. For the drug treated conditions, the drugs were added in the culture at Hour 18 and the cells were treated with the drugs for 6 hours. At Hour 24, the cells were washed with ice-cold PEG-conditioned PBS and lysed with lysing buffer. The protein was clarified by 1 min high speed centrifuging (10,000 RPM) at 4°C, aliquoted in pre-chilled microcentrifuge tubes, and then snap-frozen in liquid nitrogen and stored at −80°C. To prevent extensive hydrolysis of the active (GTP-bound) G-proteins, only one dish of cells was handled each time. The lysing procedure for each dish was done within 10 min. A small aliquot of cell lysate was later thawed and used for total protein quantification following the “BCA Protein Assay Kit” manual (Thermo Scientific). The same quantity (60 μL) of total protein from each condition at the concentration of 0.5 mg/mL was loaded in duplicate or triplicate wells of the GLISA plate. The procedures of protein binding, antigen presenting, conjugation of primary and secondary antibodies, and signal detection were conducted strictly following the manufacturer’s manual. The signal in each well was read by a microplate spectrophotometer (Molecular Devices) at the absorbance of 490 nm.

### Rheological characterizations on PA and collagen gels

An Anton Parr Shear Stress Rheometer was used for the mechanical characterization of collagen gels with the 25-mm parallel-plate geometry and a 500µm gap. A 25-mm no. 1 cover glass (VWR) was used for the top plate and a 40 mm cover glass (Fisherbrand) as the bottom plate. Both silanized cover glasses were attached to each plate of the rheometer with double-sided tape (3M 666). Polyacrylamide was deposited onto the bottom plate of the rheometer immediately before gelation, and the top plate was lowered rapidly so that the gel formed a uniform disk between the two plates. Approximately, 300 µL of solution was pipetted onto the rheometer, the gap was set to 500µm, and the sample was kept in a custom-made humidity chamber to prevent evaporation. Polymerization progress was monitored by imposing three cycles of 0.5% strain every 5 min, measuring the shear storage modulus G′ as a function of polymerization time. For strain sweep measurements, gels were subjected to 5 oscillations at 0.1 Hz at increasing amplitudes from 2 to 12% in 2% increments and 12 to 100% in 4% increments. Logarithmic frequency sweeps (0.01~3 Hz) were performed on the soft PA gel and the stiff PA gel at a 5% strain.

### Sample preparation for scanning electron microscopy (SEM)

Spreading cells on the collagen cushions were fixed with 2.5% glutaraldehyde (Polysicences, Inc.) diluted in PEG-conditioned PBS overnight at 4°C. The fixed samples were washed three times and then treated with 1% OsO_4_ in PBS for 1 hour. The samples were then thoroughly washed in deionized water three times and gradually dehydrated in ethanol with an incrementally increased concentration: 30%, 50%, 70%, 80%, 90%, and 95%. Samples were treated for at least 5 min in each ethanol bath. The samples were then soaked in fresh 100% ethanol for 5 min three times, followed by 50% ethanol plus 50% hexamethyldisilazane (HMDS) for 30 min and 100% HMDS for 30 min. After the removal of excessive HMDS, the fully dehydrated samples were air-dried in the chemical hood overnight. The samples were kept dry in a desiccator before SEM imaging. Samples were then mounted on a support with carbon tape (3M) and then covered with an 8nm layer of iridium with a sputter coater.

### Immunofluorescence staining

Cells were washed with PEG-conditioned PBS and then fixed with 4% paraformaldehyde (PFA) (Santa Cruz) for 15 min at room temperature (RT). PFA was also accordingly conditioned with PEG300 to minimize the potential osmolarity shock during fixation. After washing the cells three times with PBS, the cells were permeabilized with 0.3% Triton X-100 in PBS for 15 min and then blocked in 1% bovine serum albumin (BSA) in PBS for 1 hour at RT. After blocking, samples were incubated with the primary antibodies in PBS overnight at 4°C. Followed by a thorough wash three times in PBS for 1 hour, the samples were stained with the secondary antibodies and rhodamine-phalloidin for F-actin in the dark for 1 hour at RT. After a thorough wash in PBS three times for 1 hour, the cells were then stained with nucleus dye Hoechst before the imaging. The primary antibodies used in this study are: anti-YAP (sc-101199, Santa Cruz, 1:200), anti-vinculin (sc-73614, Santa Cruz, 1:200), anti-Myosin light chain (phospho S20) (ab2480, Abcam, 1:200), anti-VASP (sc-46668, Santa Cruz, 1:200), anti-MYO10 (PAB21665, Abnova, 1:200), and a dye-conjugated antibody anti-hemagglutinin (HA) tag-Dylight 488 (26183-D488, Invitrogen). The secondary antibodies used in this study are: Alexa Fluor 488 Goat Anti-Mouse IgG antibody (Invitrogen, A-11001) and Alexa Fluor 647 Goat Anti-Rabbit IgG antibody (Invitrogen, A-21244). We use Hoechst 33342 (Invitrogen, H3570, 1:2000) to stain the cell nuclei, and Rhodamine Phalloidin (Abcam, ab235138, 1:1000) or Alexa Flr 488 Phalloidin (Invitrogen, A12379, 1:500) to stain for F-actin.

### Drug treatments

Small molecule drugs used in this study are: ROCK inhibitor Y27632 (Cayman; 1μM and 25 μM for SNU475 cells; 5μM for HUVECs), Rac1 inhibitor NSC23766 (Tocris; 50μM), actin polymerization inhibitor Cytochalasin D (Abcam; 20 μM), Rho activator CN03 (Cytoskeleton, Inc.; 1μg/mL), Small GTPases (Rho/Rac/Cdc42) activator CN04 (Cytoskeleton, Inc.; 1μg/mL).

### Collagen compaction assay

24-well plates were coated with 500 μL 3% BSA in PBS per well at RT for 1 hour, then rinsed with PBS, and air-dried in a sterile environment. 500 μL neutralized 2mg/mL collagen mixed with SNU475 cells at densities of 0.3×10^6^ cells/mL, 1.5×10^6^ cells/mL, or 3×10^6^ cells/mL was dispensed in each well and quickly transferred in the cell culture incubators (37°C) and allowed to polymerize for 1 hour. 1mL of isotonic or PEG-conditioned cell culture medium was gently added to each well from the side to dislodge the gel. The gels were cultured for up to 48 hours and the media were changed every day for the density of 0.3×10^6^ cells/mL and twice a day for gels with higher seeding density. Macrographs were taken to record the gel area in each well over time. To estimate the plastic compaction of the gels, the cellular force was completely removed in 2% Triton X-100 by replacing 500 μL culture medium with 500 μL 4% Triton X-100 in PBS at Hour 48 followed by incubation at 4°C overnight. We define the initial gel area at Hour 0 is *A_0_*, the gel area at Hour 48 before decellularization is *A_plastic+elastic_*, and the relaxed gel area after decellularizing is *A_plastic_*.The mechanics of collagen compaction and the contribution of cellular force are evaluated as percentages defined as total compacted area *C_total_*, plastically compacted area *C_plastic_*, and elastically recoilled area normalized by plastically compacted area C*_recoilled_*:

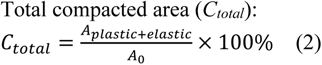

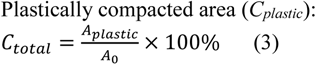

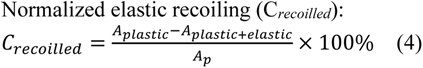

### Tube formation assay for HUVECs

Ice cold 10μL of growth factor reduced Matrigel (CORNING) was dispensed in each well of a 15-well µ-Slide (ibidi) and then gelled at 37°C for 30 minutes. After the full gelation of the Matrigel, 50 μL of endothelial cell growth medium conditioned with PEG-300 or ROCK inhibitor Y27632 with 10,000 cells was evenly plated on the Matrigel surfaces. The cells were allowed to spreading and form 2D tubular structures for 24 hours. The live GFP-labelled cells in each well were then scanned by an EVOS imaging system (Invitrogen). The cells were also fixed and their F-actin was stained by rhodamine phalloidin.

### Cell transfection

#### RacV12

The plasmid of constitutively active Rac1 (RacV12) was a kind gift from Dr. Martin Schwartz at Yale University^52^. SNU475 cells were plated in a 6-well plate at a density of 5×10^4^ cells/well and allowed to attach and proliferate for 2 days to reach a ~80% confluency. The cells were then transfected with RacV12 (0.17 μg/well) in Opti-MEM (Gibco) containing 5.5 μL Lipofectamine 3000 (Invitrogen), following the manufacturer’s protocol. The “Control” cells were the cells treated with Lipofectamine 3000 without the plasmid. After 4 hours of culture in the transfection reagent, the media were replaced with fresh complete RPMI medium and the cells were allowed to recover and express RacV12 for 2 days before using the cells for experiments. RacV12-positive cells co-expressed Hemagglutinin (HA) which was determined after fixation by immunofluorescence staining with the antibody HA-488. The baseline signal for RacV12-positive cells was defined as higher than the maximum mean intensity of the 488 signal of the Lipofectamine-treated “control” cells.

#### LifeAct-RFP

SNU475 cells in a well of a 24 well plate (2×10^4^ cells/well) were treated in 500 μL complete RPMI medium with 8 μg/mL polybrene (Sigma) and 2μL (2 multiplicity of infection) rLV^Ubi^-LifeAct-TagRFP lentiviral vector (ibidi) for 16 hours, The medium was then replaced with fresh medium and cells were allowed to expand for 2 days. Stably transfected (RFP positive) cells were then selected with 2 μg/mL puromycin (Gibco) in the culture medium.

### Osmotic gradient device

A PDMS sheet with a 2mm thickness was cut to fit on a round glass cover slip (diameter: 40mm). A ~4mm width, 13mm long channel was cut out of the PDMS block by a scalpel blade. A reservoir was then cut out on each side of the channel by an 8mm biopsy punch twice so that each reservoir can roughly contain 200 μL culture medium. The PDMS block and the glass cover slip were permanently bond together after a surface treatment in an air plasma cleaner. This open channel device was sterilized in 70% ethanol and then air dried in a tissue culture cabinet. Two small pieces of PDMS was used to plug the two ends of the channel to create a small vessel. Approximately 120 μL collagen (2mg/mL) with a cell density of 10,000 cells/mL (SNU475 cells) was then filled up the straight channel without leakage into the reservoirs. After gelling the collagen at 37°C for 1 hour, the PDMS plugs were removed and ~ 200 μL isotonic medium was added in both reservoirs. The medium was allowed to diffuse into the gel at 37°C for 1 hour from both sides until the color of the gel became uniform. Then media were changed to fresh isotonic medium and 4% PEG medium respectively in the two reservoirs. The device was carefully transferred into the incubator and cultured for 24 hours. The same medium type was replenished at Hour 12. At Hour 24, the gel on the device was fixed and stained with rhodamine phalloidin and Hoechst. A confocal image volume stack was taken to cover the whole length of the channel with a 1200 μm width and a 1000 μm depth.

### Tumor invasion in the 3D collagen ECM

SNU475 tumor spheroids were made from 1,000 cells in 100uL culture medium per well in an ultra-low attachment 96 well round bottom plate (CORNING) for 2 days. The spheroids were collected and mixed into a 2mg/mL neutralized rat tail collagen precursor solution. It was made sure about 4-5 spheroids were incorporated in 60uL gel solution per well in dopamine-HCl functionalized glass bottom 12-well plates (Cellvis). The gel droplets formed domes and were allowed to solidify for 1 hour at 37°C, followed by the addition of isotonic or PEG-conditioned media with or without drug treatments. The spheroids were then allowed to grow for 48 hours.

### Applying mechanical compression on cells

Planar mechanical compression was applied on cells, following the previously reported method^65^. In brief, LifeAct-SNU475 cells (8,000 cells/well) were plated on 50μg/mL collagen I coated 0.4 μm pore polyester transwells in 6-well plates (Corning). Round agarose cushions (diameter: 20mm) were cut out from a ~2mm thick sterile 0.8% (w/v) agarose sheet formed in a petri dish. The agarose cushions were gently placed on top of the cell covered transmembranes after the cells were allowed to attach for 2 hours. The cells were considered under minimum compression from the agarose cushions due to the buoyancy. For the compressed condition, a small cup filled with stainless steel balls weighing 25 g in total was gently placed on top of the agarose cushion, which introduced approximately 5.8 mmHg (~ 773Pa) pressure on the cells. Both sides of the transwells were filled with complete culture medium. The cells in this experiment setup were cultured for 24 hours. The cells were then washed with PBS and fixed with 4% PFA before the weights and cushions were removed. The transwells were transferred to a glass bottom dish for confocal imaging.

### Schematic illustrations

Schematics in the manuscript were designed and drawn using an open-source software Inkscape.

### Microscopic imaging and data analysis

All fluorescent imaging was conducted on a confocal microscope (Leica SP8) equipped with an environmental chamber (37°C, 5% CO_2_) for the live-cell imaging. For the time-lapse imaging of cell spreading, long-distance 10x or 20x objectives were used. Fluorescent images were acquired with the long-distance 20x objective or a water immersion 40x objective. For live cell imaging and, cells were stained with CellTracker Green (Invitrogen, C7025, 1:1000 dilution) for 25 min, washed with PBS, centrifuged, and resuspended in a fresh culture medium. For the high-resolution cell volume reconstruction and calculation, live CellTracker-labelled cells were imaged with the water immersion 40x objective with a z-step of 0.1 μm. All image analysis was performed with Fiji (ImageJ)^64^. Cell spreading area and shape descriptors (solidity and circularity) were quantified by manual tracing or generating a binary mask from the fluorescent images. FA size was characterized based on the confocal fluorescent images (max projection), using ImageJ with this sequence: “Subtract Background>Gaussian Blur, sigma=0.6 >Threshold>Convert to Mask>Analyze Particles”. YAP subcellular localization was determined by the ratio of YAP mean intensity in the nuclei and cytoplasm following the quantifying method published previously^28,29^. Masks of the nucleus or whole cell shape were manually traced or segmented with ImageJ. Using the high resolution live CellTracker images and ImageJ’s “3D Objects Counter”, we estimated cell volume by the voxel number from the thresholded CellTracker signal, following a previously reported protocol^3^.

### Statistics

The statistical tests, the number of biological replicates, and technical replicates were stated in figure legends. The statistical comparisons were performed using either Origin 2020b (OriginLab) or GraphPad Prism 9 (GraphPad).

---

**Supplementary Figure 1.**
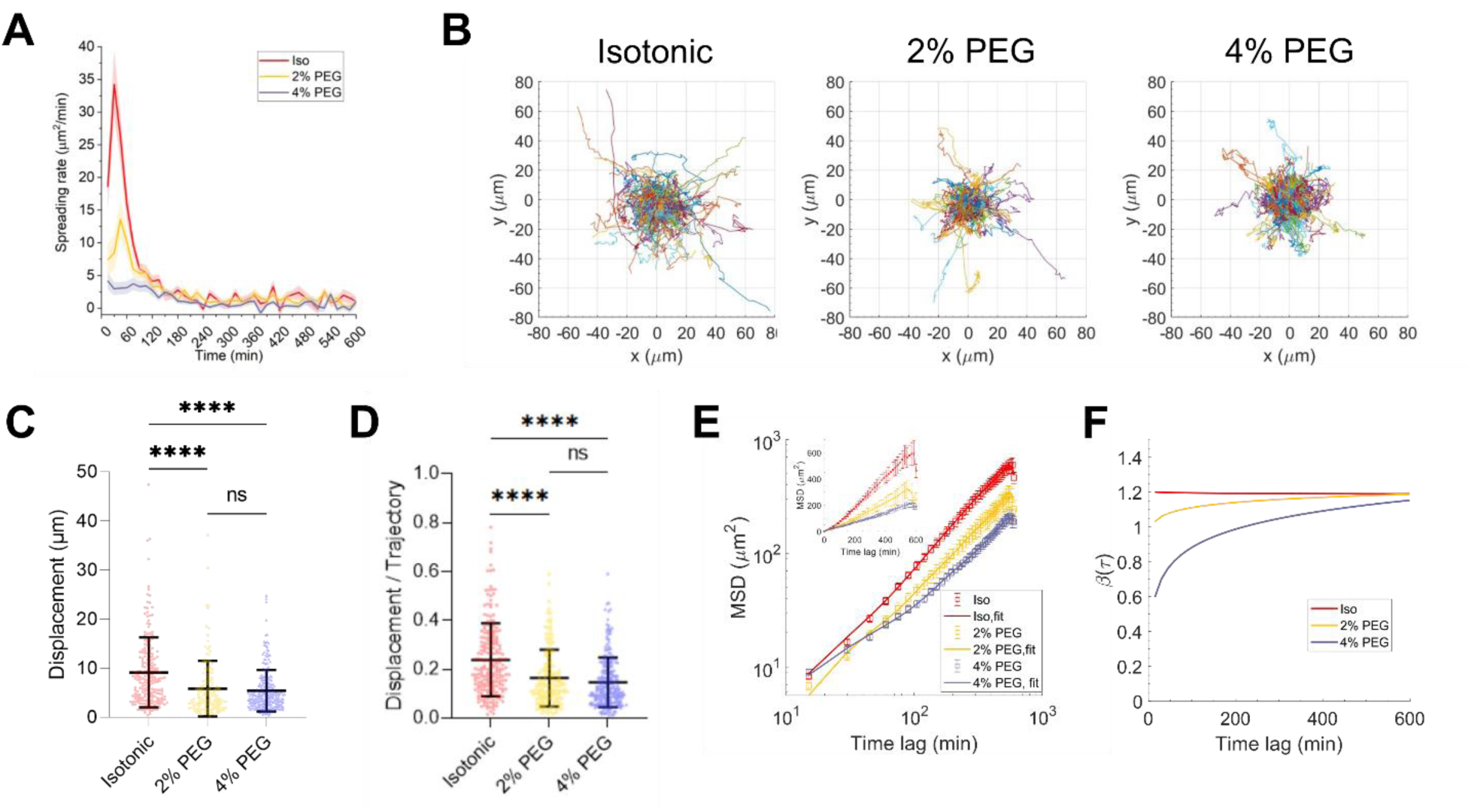
Characterization of spreading dynamic of cells under compression. (A) Spreading rate of the cells between 15-min time intervals over the course of the 10-hour time-lapse imaging. In the isotonic condition and 2% PEG condition, the cells rapidly spread in the first hour and then continue spreading at a slowed-down rate afterward. The 4% PEG renders a rather stably low spreading rate. Overall, the non-compressed cells exhibit a much higher spreading rate. Data represent mean±s.e.m. (B) Cell centroid tracking during cell spreading shows: (C) less movement and (D) less directed motion of the compressed cells, which is further reflected in the mean square displacement (MSD) plot (E). n= 209, 150, and 247 cells for isotonic, 2% PEG, and 4% PEG conditions respectively, from one experiment. In the log scale, the mean trajectory from each condition is fitted to a polynomial curve shown as the solid lines. (F) The logarithmic derivative β(*t*) of the MSD trajectories characterizes the cell movement as the anomalous diffusion model. The compressed cells have lower β values, indicating a more “confined” and subdiffusive motion. The inset in (E) shows the data in the linear scale. In (C) and (D), data represent mean±s.d. Significant comparisons were determined with a one-way ANOVA with a Tukey post hoc test. (*p<0.05; **p<0.01; ***p<0.001; ****p<0.0001). In (E), the data are shown as mean±s.e.m.

**Supplementary Figure 2.**
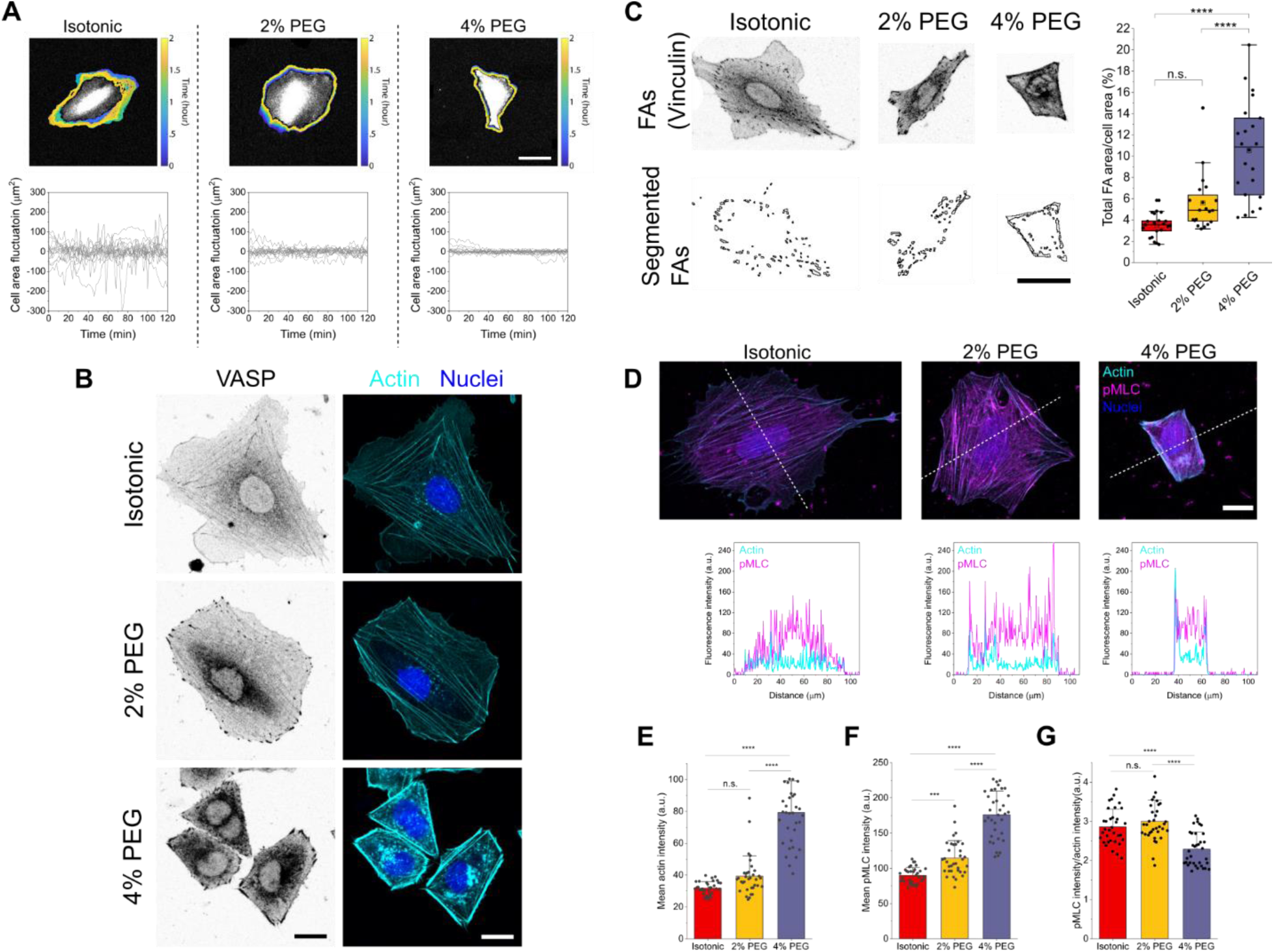
Characterizing adhesion-actin machinery of volumetrically compressed cells. (A) Cell area fluctuation of fully attached cells over 120 min. The area fluctuation around zero over time is the cell area subtracted by the mean cell area. Scale bar: 50 μm. n=15 cells from three independent experiments for each condition. (B) Fluorescent images (inverted intensity) showing VASP puncta accumulated at the end of peripheral actin bundles around the cell boundary under compression. Scale bar, 20μm. (C) Fluorescent images with inverted intensity showing vinculin puncta and representative FA segmentation. This was used to estimate the ratio of FA in the total cell area of individual cells. N=21, 20, and 22 cells respectively from two independent experiments. Scale bar, 50 μm. (D) Fluorescent images showing the co-localization of F-actin and pMLC. Scale bar, 20 μm. The actin intensity, pMLC intensity, and pMLC intensity normalized by the actin intensity in the cells under different conditions are shown in (E), (F), and (G) respectively. Statistic analyses in (C,E,F,G) were calculated by one-way ANOVA with a Tukey multiple comparisons test (*p<0.05; **p<0.01; ***p<0.001; ****p<0.0001; ns: not significant).

**Supplementary Figure 3.**
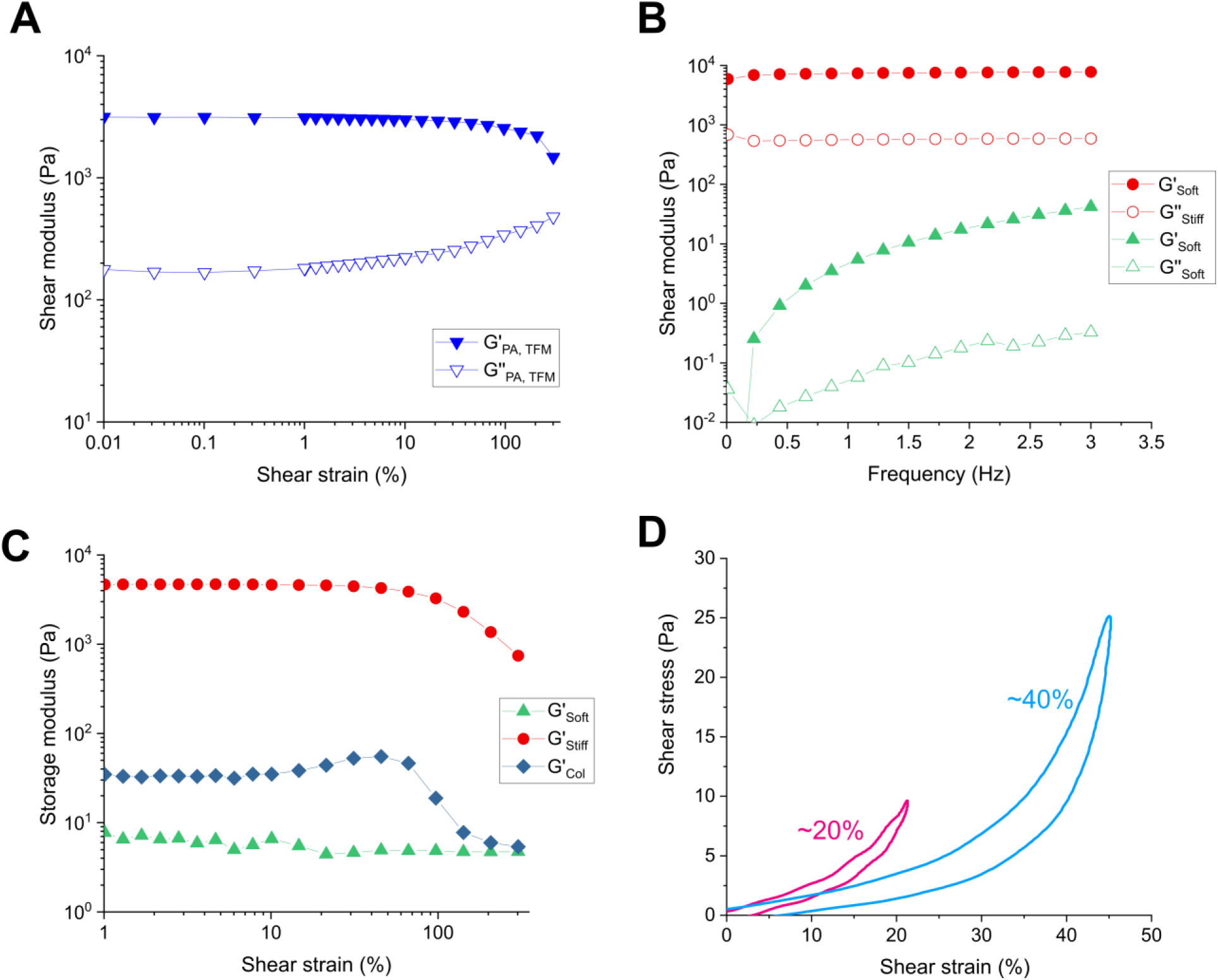
Rheological characterization of polyacrylamide (PA) gels and collagen. (A) A strain sweep (1%-300%) was used to characterize the mechanical property (shear storage modulus: G’; shear loss modulus: G’’) of the PA gel (with microbeads) used for the traction force microscopy. The shear storage modulus (G’) of the gel substrate at 1 Hz is ~ 3 kPa, therefore the estimated Young’s modulus is ~9 kPa, assuming PA is nearly incompressible (Poisson ratio equals 0.5). This Young’s modulus was used to calculate the strain energy generated by the cells in Fig 3. (B) Frequency sweeps (0.01~3 Hz) were performed on the soft PA gel and the stiff PA gel at a 5% strain. The “soft” and “stiff” PA gels were used to demonstrate cell spreading on an elastic substrate in Fig 4. (C) Strain sweeps (1%-300%) at the frequency of 1 Hz were also performed to characterize both soft and stiff PA gels, as well as the collagen gel (2mg/mL). With the strain increasing, the PA gels exhibit constant modulus (elastic behavior), while collagen exhibit strain stiffening behavior before failure. The shear storage moduli of the soft gel, the stiff gel, and collagen are 5.5 Pa, 7.5 kPa, and 33 Pa at 1 Hz, respectively. (D) Cyclic shear rheology reveals a nonlinear viscoplastic behavior of collagen at 20% and 40% strains. The area discrepancy under the loading and unloading curves of each cycle indicates the plasticity of collagen.

**Supplementary Figure 4.**
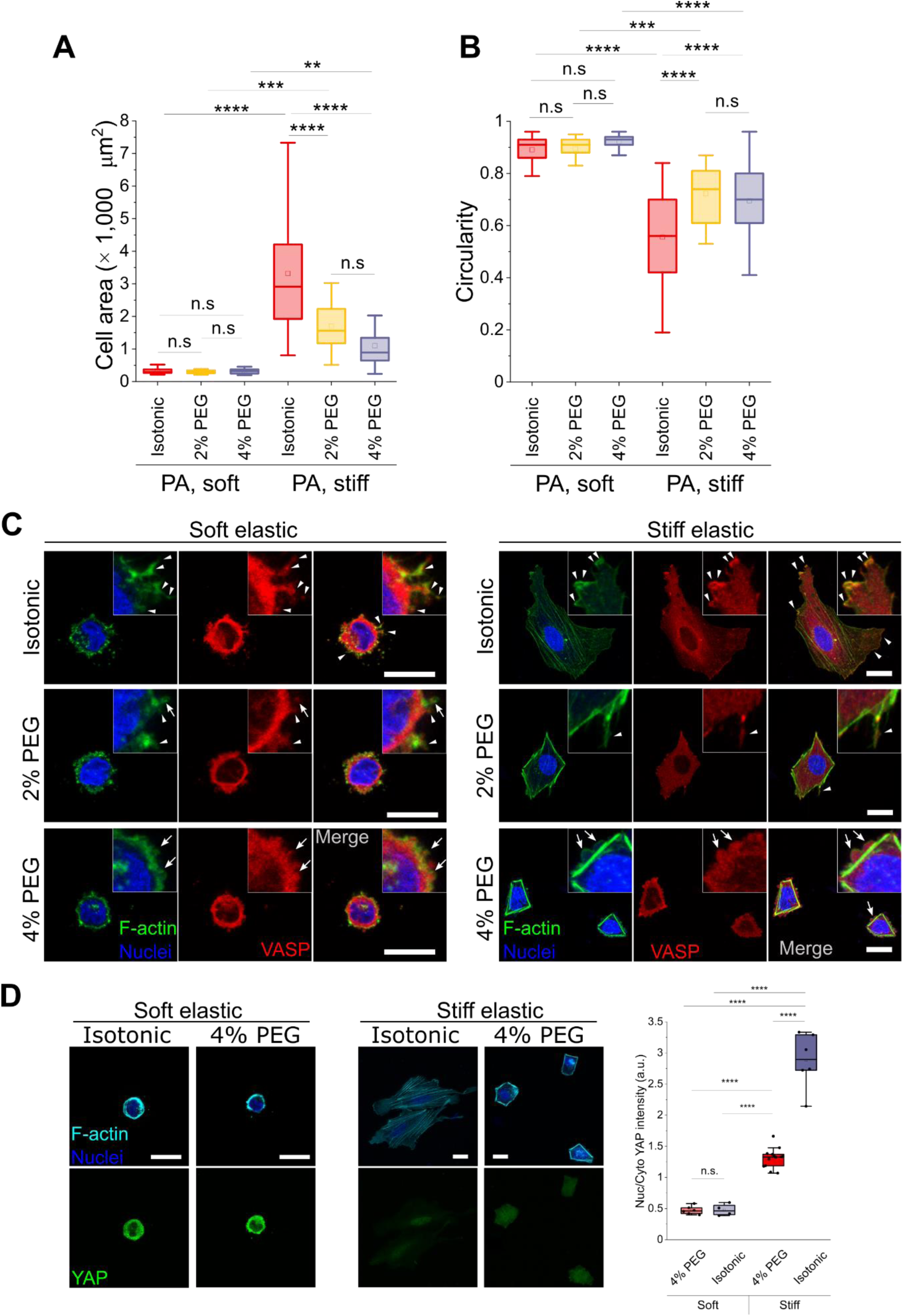
Mechanosensing of cells on the soft and stiff elastic substrate under compression. (A) Cell area and (B) cell circularity measurements of non-compressed and 2% PEG or 4% PEG compressed cells seeded on the soft elastic, collagen, and stiff elastic substrates. (C) Fluorescent images of VASP and F-actin of the cells on soft and stiff elastic substrates under varied compression. The close-up images show the distinct morphologies of cell actin processes with VASP. Arrowheads indicate filopodia/lamellipodia-like protrusions and arrows indicate blebbing. (D) Fluorescent images of YAP and F-actin of the cells on soft and stiff elastic substrates under isotonic and 4% PEG conditions. YAP nuclear translocation under varied compressions was quantified. Scale bars, 25 μm.

**Supplementary Figure 5.**
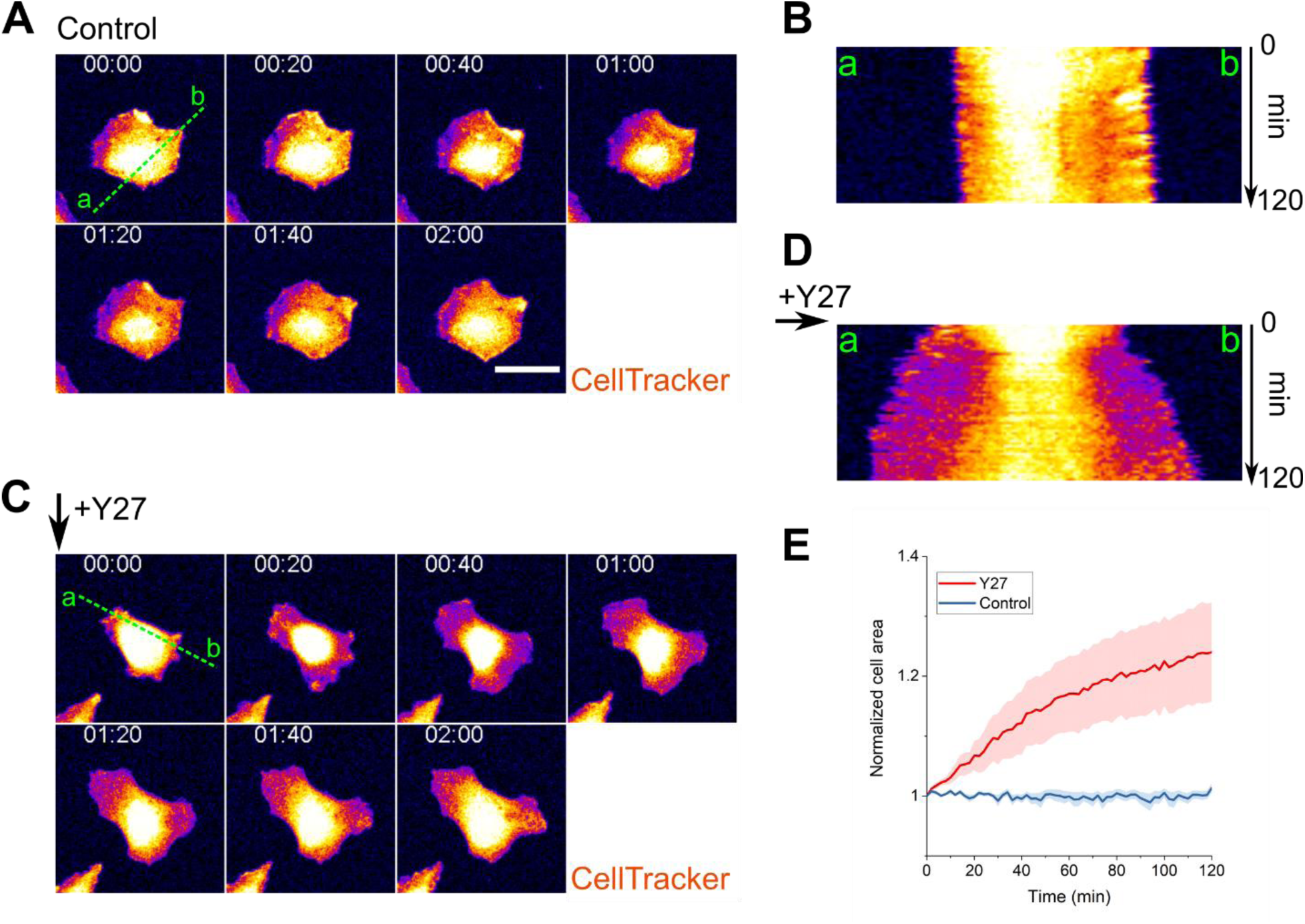
High dosing of ROCK inhibition significantly recovers cell spreading area on 2D under compression. (A) Time-lapse imaging (2 hours) of a fully spreading 2% PEG compressed cell with minimum area change and edge ruffling shown by the kymograph in (B). (C) Time-lapse imaging (2 hours) showing significant cell area recovery of a 2% PEG compressed cell upon the treatment of 25 μM ROCK inhibitor (Y27632). The expansion of the edge across the green dotted line is shown by the kymograph in (D). (E) Characterization of the change in cell area over 2 hours with and without ROCK inhibition under 2% PEG compression, normalized by the area at time 0. n=9 cells for Control, and 8 cells for the Y27 treatment. Scale bars, 50 μm. Representative imaging is shown in Supplementary Video 5.

**Supplementary Figure 6.**
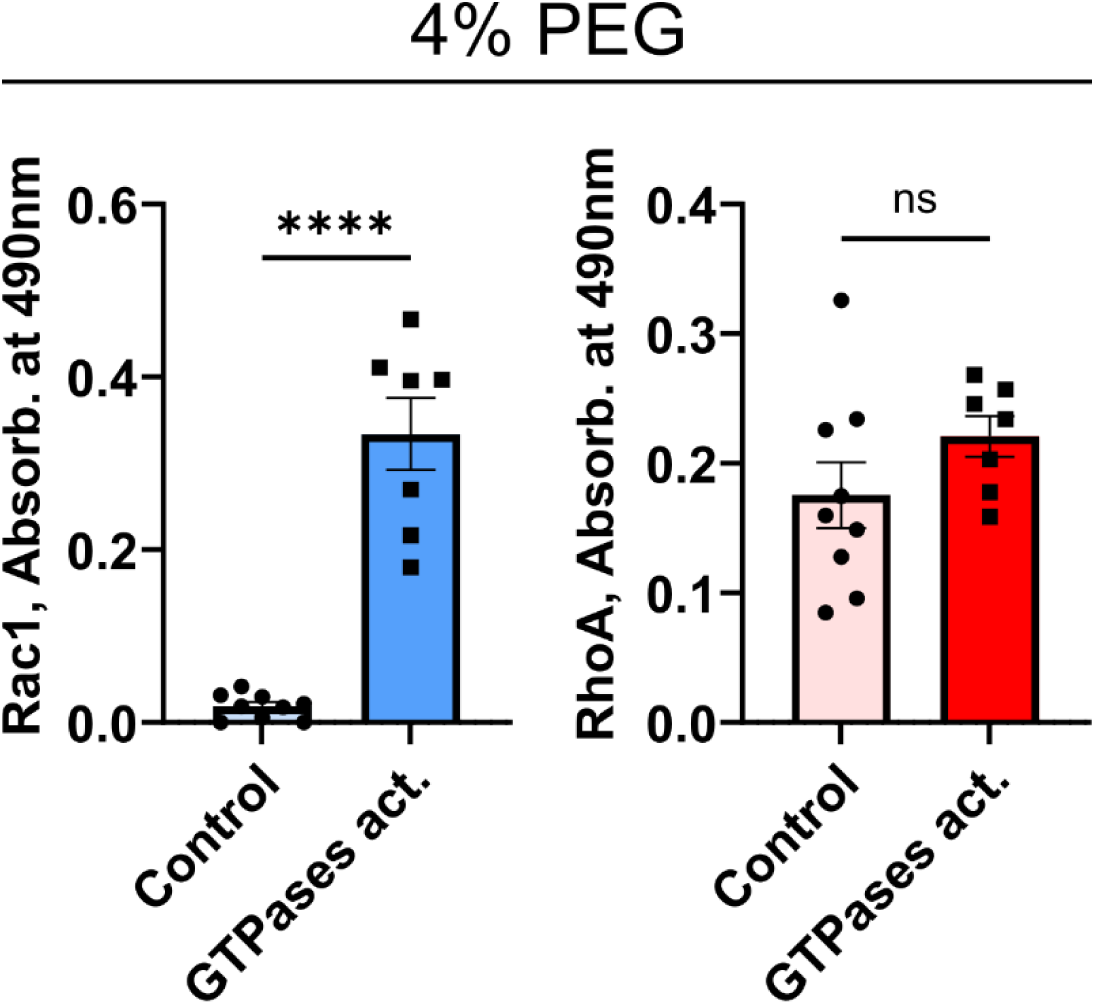
Effects of GTPases activator (CN04) on Rac1 and RhoA activities. GLISA assays were used to determine levels of Rac1 and RhoA activities after 6-hour treatment of GTPases activator CN04 (GTPases act.). This activator shows a long-lasting effect on Rac1 activation in 4% PEG-compressed SNU475 cells but not on the RhoA. Plots are shown as Mean±s.e.m. n=9 replicates for Control, and 7 replicates for GTPase activator, from three independent experiments. Two-tailed t-tests are used to determine the statistical difference (****p<0.0001; ns: not significant).

**Supplementary Figure 7.**
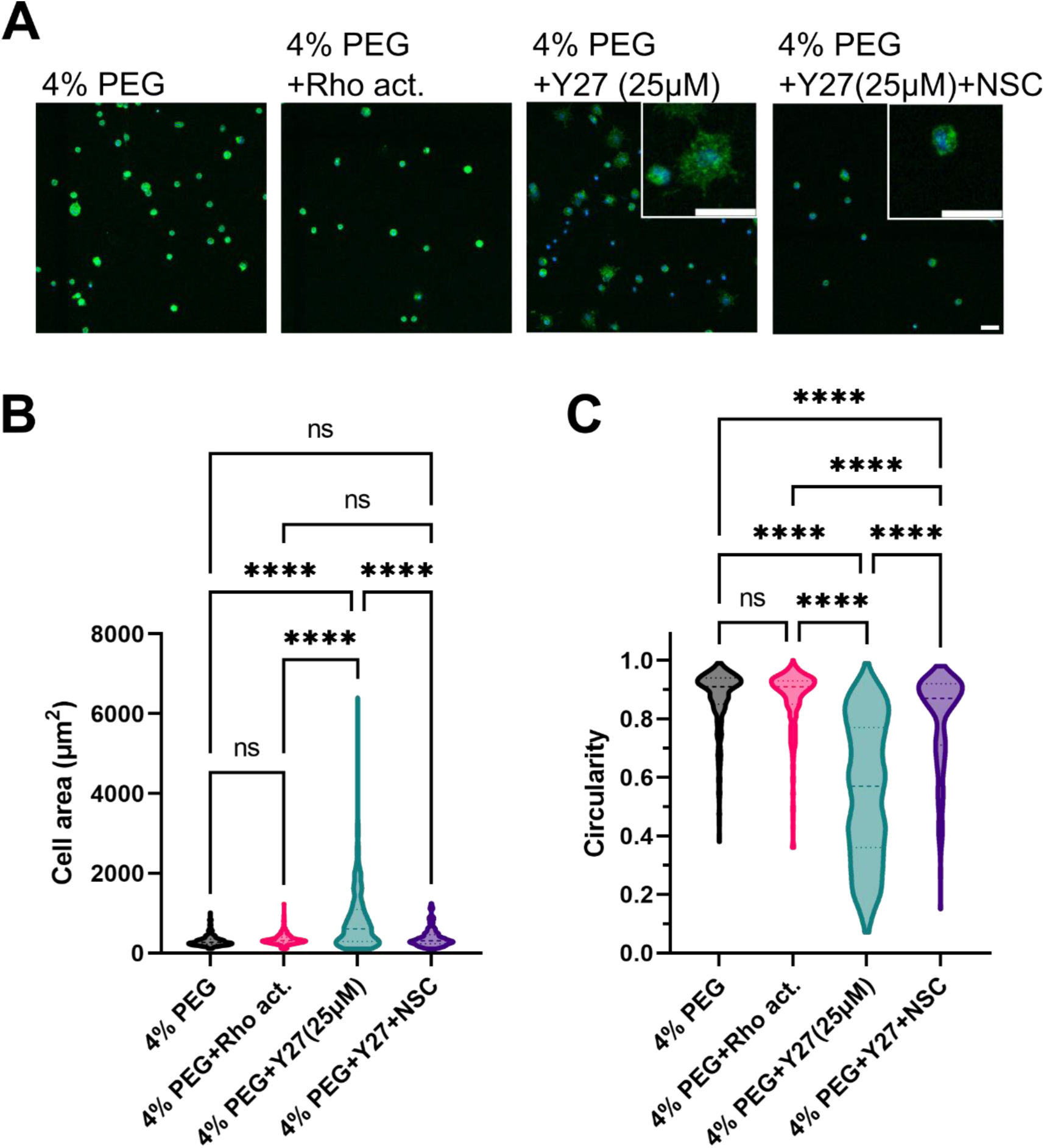
Characterizing cell spreading of 4% PEG compressed cells on soft collagen cushions with pretreatments of small molecule drugs. (A) Representative images (green: actin; blue: nuclei) of the cells on soft collagen cushions 24 hours after cell seeding. These cells are pretreated by Rho activator (CN03), ROCK inhibitor Y27632 (25μM), or Rac1 inhibitor NSC23766, and stay in the same drug treatments throughout the spreading. Scale bars, 50 μm. Cell morphology is characterized by cell spreading area (B) and circularity (C). N=303, 316, 359, 203 cells for the corresponding conditions from two independent experiments. Statistic significance was calculated by one-way ANOVA with a Tukey multiple comparisons test (****p<0.0001; ns: not significant). We demonstrate that solely activating Rho does not rescue cell spreading. Relaxing cell contractility significantly rescues the compression-suppressed cell spreading and protrusion formation, which can be inhibited by the Rac1 inhibitor. This demonstrates the inhibitory effects of Rho/ROCK contractility on Rac1-guided protrusion formation.

**Supplementary Figure 8.**
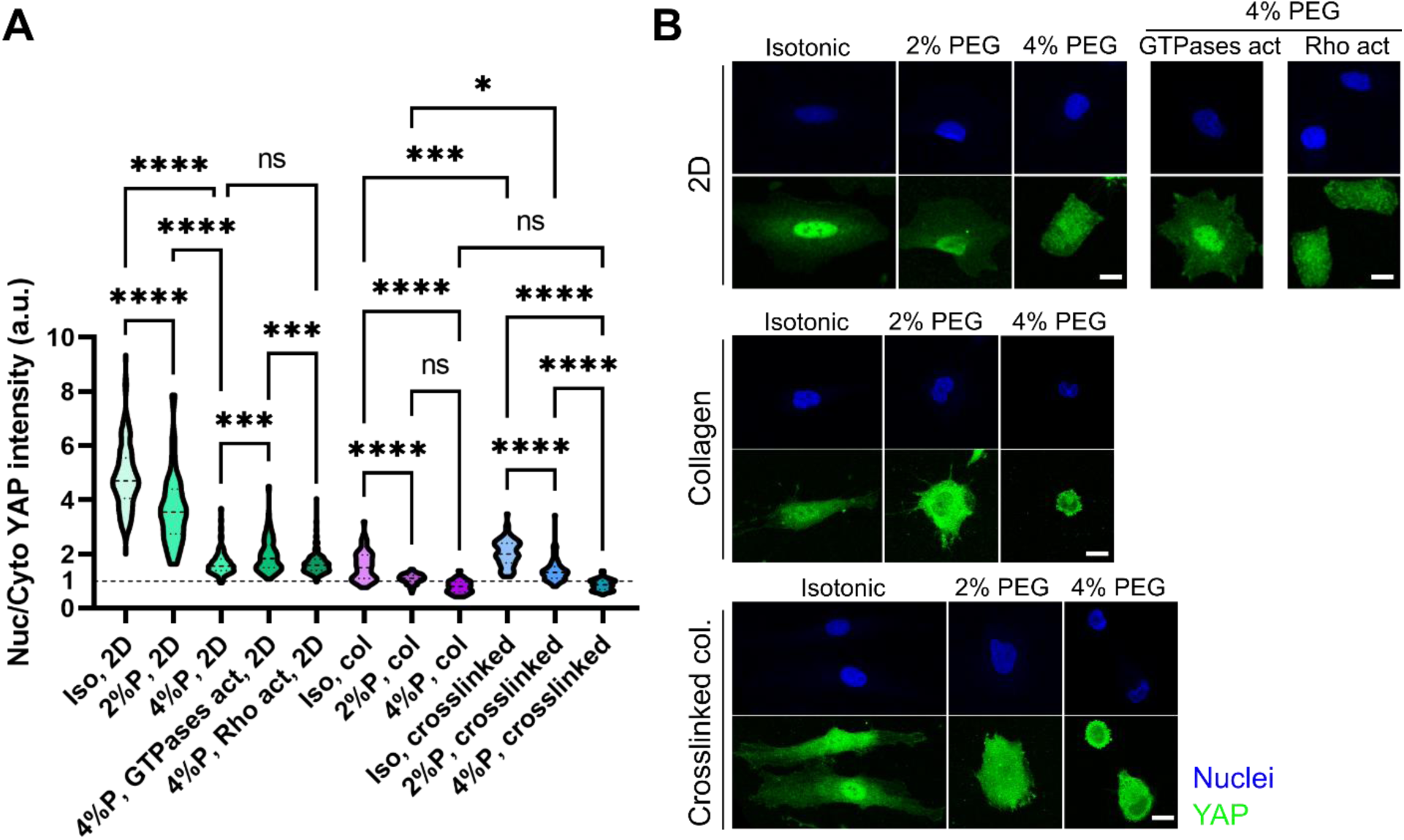
The interplay between compression and substrate type determines cell spreading and YAP subcellular localization. (A) Characterization of YAP nuclear translocation cross compression conditions (Iso: isotonic; 2%P: 2% PEG; 4%P: 4% PEG), drug treatments, and substrate types (2D: collagen coated rigid surface; col: collagen cushion; crosslinked: glycated collagen cushion). n=129, 32, 186, 140, 216, 59, 125, 43, 75, 98, 120 cells for the corresponding conditions from two to three independent experiments. Statistic significance was calculated by one-way ANOVA with a Tukey multiple comparisons test. (*p<0.05; **p<0.01; ***p<0.001; ****p<0.0001; ns: not significant) (B) Representative immunofluorescent images showing YAP intensity with respect to the nuclei under different conditions.

**Supplementary Figure 9.**
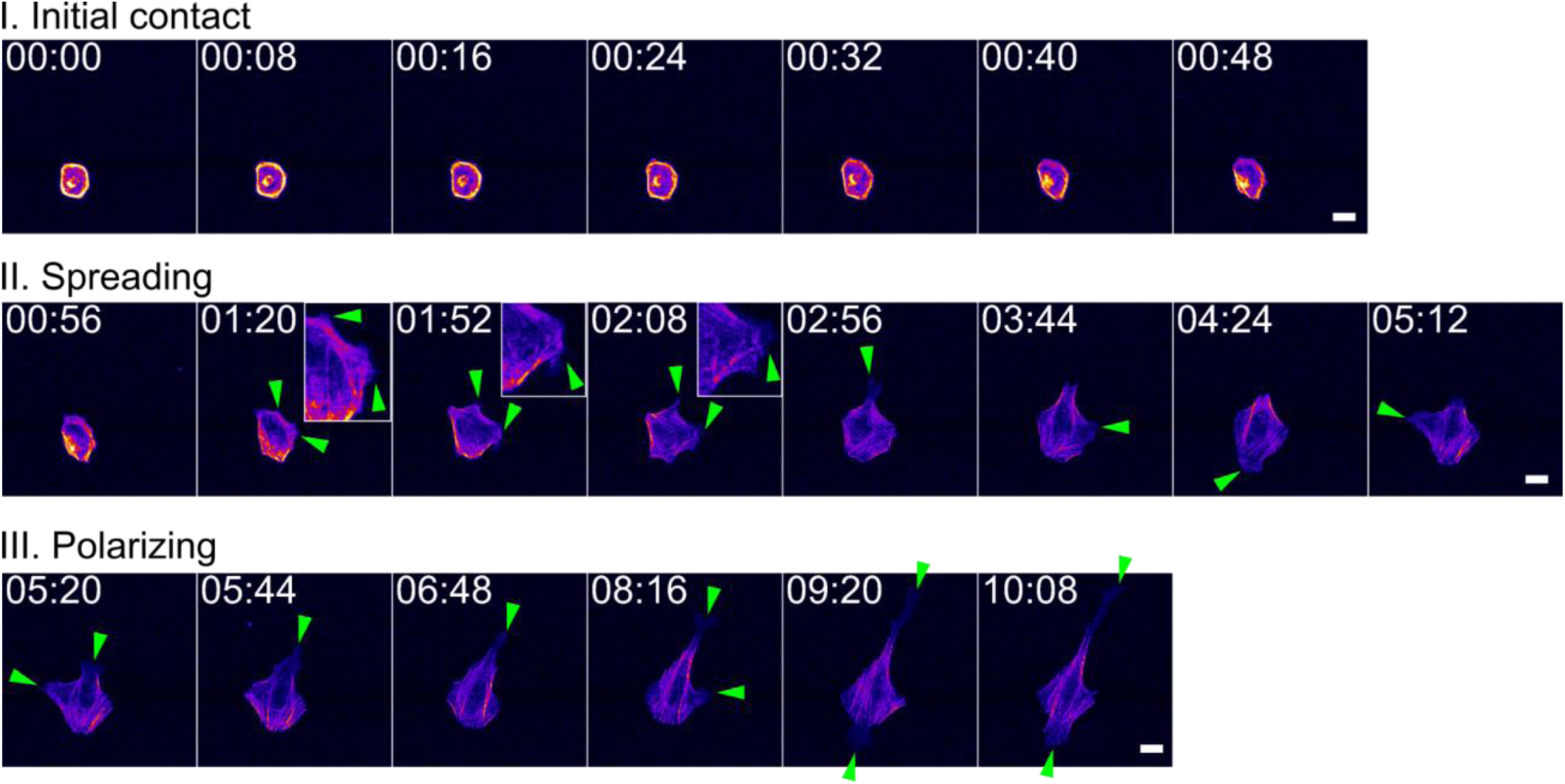
Spreading of a LifeAct-labelled SNU475 cell on the 2D rigid surface over 10 hours in the isotonic condition. There are three stages of cell spreading: (I) Initial contact: the round cell adheres to the substrate with only cortical actin. Over time, the cortical actin flattens out to become peripheral actin bundles. The bundles extend the cell body and establish multiple adherent vertices with no evident lamellipodia formed. At the moment, the cell exhibits a distinct polygonal shape. (II) Spreading: sheet-like dynamic lamellipodia are formed from the vertices (enlarged in the insets) to facilitate the separation of the peripheral actin bundles into stress fibers. (III) When reaching an optimal area, the cell shifts into a more polarized shape by generating protrusions in two opposing directions. Green arrows indicate the dynamic protrusions guiding the cell spreading and polarizing. Scale bars, 20 μm. Representative imaging is shown in Supplementary Video 12.

**Supplementary Figure 10.**
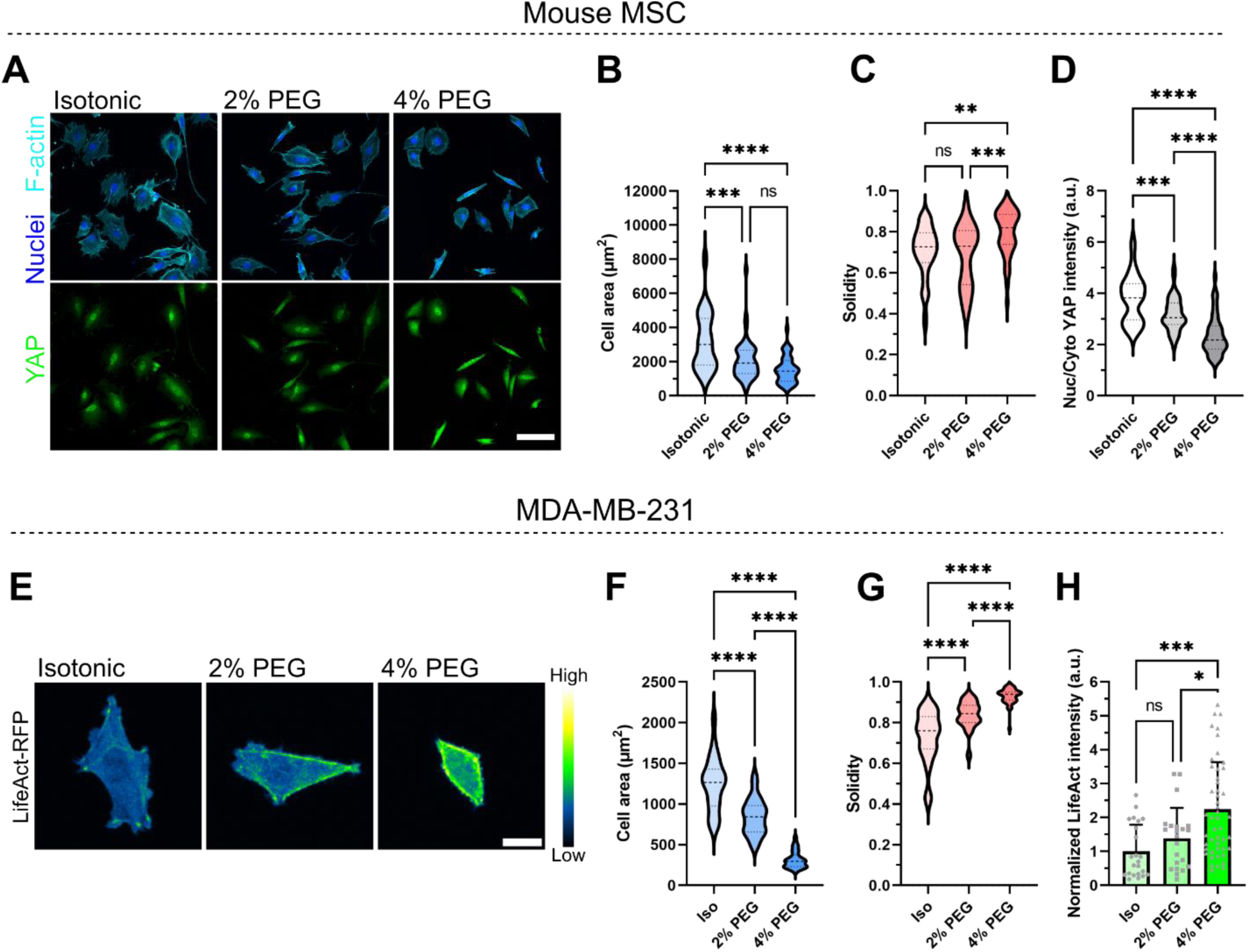
Volumetric compression regulates dynamic cells’ spreading, cytoskeleton arrangement, and YAP translocation. (A) Mouse mesenchymal stem cells (MSC) exhibit distinct spreading morphology and actin arrangement under volumetric compression in a PEG concentration-dependent manner. Scale bar, 100 μm. Volumetric compression reduces (B) cell spreading area, (C) increases cell shape solidity, and (D) reduces YAP nuclear translocation. (E) Actin arrangement and intensity of metastatic breast cancer cells MDA-MB-231 labeled by LifeAct-RFP. Scale bar, 20 μm. Consistent with liver cancer cells SNU475, this cancer cell line under compression also exhibits (F) reduced spreading area, (G) increased solidity, and (H) higher actin intensity. MSC: n= 45, 44, 49 cells for the corresponding condition; MDA-MB-231: n=23, 22, 48 cells for the corresponding condition. Statistic significance was calculated by one-way ANOVA with a Tukey multiple comparisons test. (*p<0.05; **p<0.01; ***p<0.001; ****p<0.0001; ns: not significant)

**Supplementary Figure 11.**
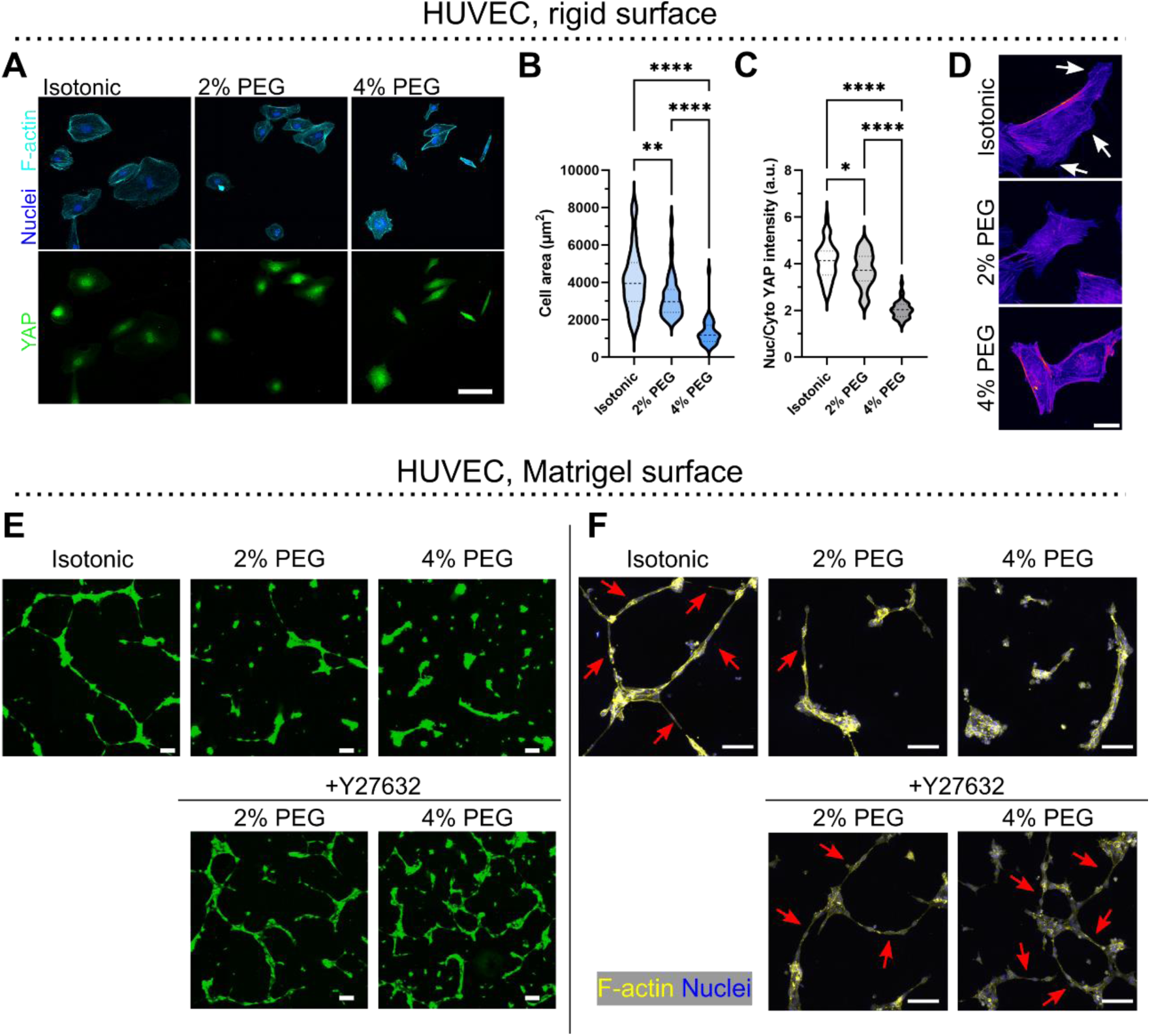
Volumetric compression regulates HUVEC spreading, actin protrusions, and network formation. (A) Human umbilical vein endothelial cells (HUVECs) respond to volumetric compression in 2D culture with distinct actin arrangement. Scale bar, 100 μm. With the increase in compression, the cells reduce (B) spreading area and (C) YAP nuclear translocation. n=44, 45, 40 cells for the corresponding conditions. Statistic significance was calculated by one-way ANOVA with a Tukey multiple comparisons test. (*p<0.05; **p<0.01; ***p<0.001; ****p<0.0001; ns: not significant) (D) Confocal images (intensity map) with higher magnification show the actin arrangement. Under compression, the cells lose the sheet-like lamellipodia and accumulate actin bundles at the cell boundary. The arrows indicate the lamellipodia formation of HUVECs in the isotonic condition. Scale bar, 20 μm. (E) Tube formation of GFP-labeled live HUVECs 24 hours after being plated on the basement membrane surface (Matrigel) under five conditions: isotonic, 2% PEG, 4% PEG, and the two PEG conditions with ROCK inhibitor (Y27632, 5 μM). (F) Confocal images of F-actin showing cell-cell interaction in the tube formation assays. Scale bars in (E) and (F), 100 μm. We show that the compression suppresses 2D tubular formation by inhibiting cell extension and cell-cell connections. In the PEG-conditioned medium, the cells tend to aggregate locally and fail to establish long-range, connected networks. Reducing cell contractility by Rho/ROCK inhibition recovers cell extension and cell-cell connections, highlighted by the arrows in (f).

**Supplementary Figure 12.**
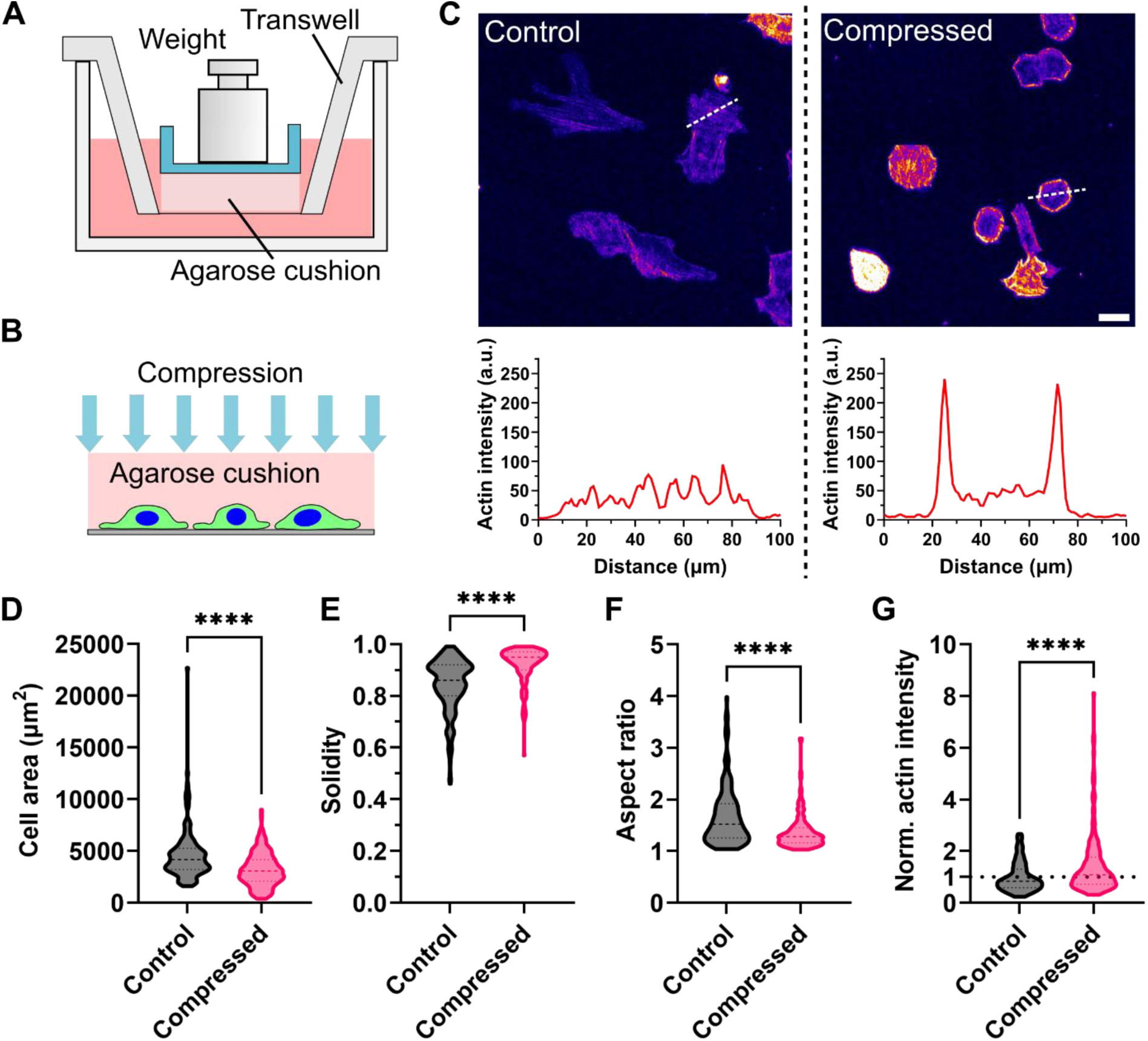
Physical compression regulates actin cytoskeleton organization and cell morphology. (A) Schematic of the experimental setup for applying mechanical compression on SNU-475 cells. 5.8mmHg (~773 Pa) pressure were applied on cells after 2-hour cell attachment on a collagen coated transmembrane. The cells were kept in cultured under compression for 24 hours. (B) Close-up schematic showing the mechanical compressive stress was applied on the cells through a soft agarose cushion. (C) Confocal imaging on LifeAct-labled actin of the non-compressed vs. compressed cells. Compared to the non-compressed cells, the compressed cells did not generate lamellipodia protrusions and exhibited smaller cell spreading area and polygonal morphology highlighted with distinct thick peripheral actin bundles. Morphology (area, solidity, and aspect ratio) and actin intensity were quantified in (D) to (G).

